# A Cross-scale Causal Mapping Framework Pinpoints Macrophage Orchestrators of Balanced Arterial Development

**DOI:** 10.1101/2025.10.08.681286

**Authors:** Jonghyeuk Han, Dasom Kong, Erica Schwarz, Felipe Takaesu, Seunguk Lee, Chae-lin Kim, Evan Yang, Minseon Park, Eunha Kim, Wook Kim, Abhay B. Ramachandra, Edward P. Manning, Jay D. Humphrey, Hyun-Ji Park, Michael E. Davis

## Abstract

Postnatal pulmonary arteries experience an abrupt surge in flow that demands tightly coordinated remodeling of vascular structure and mechanical compliance. However, the cellular programs orchestrating these multiscale adaptation remain poorly defined, partly due to the absence of analytical tools integrating gene–cell–tissue scales. Here, we present CausaLink, a cross-scale causal mapping framework that predicts how altered gene expression propagates through gene–cell–tissue networks by integrating time-course transcriptomic and tissue morpho-mechanical data. We constructed a single-cell transcriptomic atlas of 11,143 proximal pulmonary artery-derived cells from C57BL/6J mice across five developmental stages (P2, P10, P21, P42, and P84), revealing dynamic lineage transitions and transient mesenchymal population specifying into fibroblasts and smooth muscle cells from P2 to P21. Using this framework, we identified *Mgl2*⁺ macrophages as central regulators promoting lumen expansion while preventing pathological wall thinning or thickening. To validate these predictions, we established a human induced pluripotent stem cell–derived arterial assembloid enriched in MGL^high^ macrophages. In this model, MGL^high^ macrophages supported lumen enlargement while maintaining overall wall thickness and induced formation of an adventitia-like fibroblast layer that recapitulated approximately 80% of the native adventitial fraction with balanced collagen-elastin remodeling. By linking predictive modeling to human organoid validation, this study establishes a cross-scale workflow for tracing how gene programs shape vascular architecture, offering mechanistic insights and a foundation for predictive regenerative medicine.

## Main

The transition from fetal to postnatal life constitutes one of the most dramatic physiological shifts in mammalian biology. This transformation is particularly evident in the pulmonary circulation, where closure of the ductus arteriosus and foramen ovale shortly after birth redirects right ventricular output through the lungs, resulting in an eight- to tenfold increase in pulmonary arterial flow despite a concurrent decline in arterial pressure^1, 2^. This hemodynamic shift transforms the main pulmonary artery from a low-flow, high-resistance conduit into a high-flow, low-resistance circuit, demanding rapid initial remodeling followed by balanced maturation of vascular architecture^3, 4^. Precise regulation across multiple scales is essential, as failure of these adaptations can lead to persistent pulmonary hypertension of the newborn. Moreover, early disruptions in this remodeling cascade predispose to later lung dysfunction, as well as right ventricular strain or failure during adolescence or adulthood^5–7^. Despite its importance, the cellular programs orchestrating the coordinated development of vascular structure and function remain incompletely understood, which requires integrative approaches that bridge molecular, cellular, and tissue-level analyses.

Over the past two decades, transcriptomic profiling, *in vitro* mechanical and histological assessments, and *in vivo* morphometric and hemodynamic analyses in mice have advanced our understanding of vascular development by systematically cataloguing changing transcriptional profiles, cellular phenotypes, and biomechanical properties across developmental stages^8–10^. Single-cell RNA sequencing (scRNA-seq) has enabled delineation of key vascular and immune cell populations^11, 12^, while genome-wide association studies (GWAS) and transcriptome-wide association studies (TWAS) have linked gene variants to vascular traits^13, 14^. Nevertheless, these methods primarily identify statistical associations rather than causal mechanisms. Critically, these approaches have been largely restricted to mapping single gene–single trait relationships and are unable to capture how multiple tissue-level outcomes—such as vascular dimensions and compliance—co-evolve through shared regulatory programs^14–17^. Furthermore, single-cell transcriptomic analyses require destructive tissue processing, making it impossible to perform parallel histological or biomechanical analyses on the same specimen, thereby precluding direct multimodal integration^18, 19^. Consequently, such constraints limit causal modeling across biological scales.

To our knowledge, no existing method reconstructs causal hierarchies across diverse datasets obtained independently from different modalities, such as transcriptomics and biomechanics. We address this unmet need with CausaLink, a causal integration framework that infers directed gene–cell–tissue relationships from independently acquired transcriptomic and morpho-mechanical datasets, enabling predictive modeling of multicellular tissue development and adaptation. Bridging computational inference with experimental validation is essential to establish causal mechanisms; therefore, we employ a human induced pluripotent stem cell (hiPSC)-derived arterial assembloid that recapitulates native vascular architecture and remodeling dynamics. Through this framework, we identified a macrophage (Mac) subpopulation as a novel causal regulator candidate and validated its predicted role in the hiPSC-derived arterial assembloid platform. Together, our findings demonstrate an integrated computational-experimental approach toward predictive physiology and pathophysiology, offering a strategy to infer and test causal mechanisms underlying tissue development, homeostasis, and disease progression, with promise to guide rational design in therapeutics including tissue engineering and regenerative medicine.

## Results

### Cellular, Structural, and Functional Transitions during Pulmonary Artery Development

Postnatal development of the pulmonary artery is characterized by coordinated transitions in cellular phenotype, tissue architecture, and mechanical function as the vessel accommodates increasing circulatory demands^20, 21^. Yet, the cellular programs and lineage dynamics guiding this growth and remodeling remain poorly understood, particularly during the early postnatal period when structural reorganization is most rapid^22, 23^. To systematically chart these transitions, we constructed a single-cell transcriptomic atlas spanning five developmental stages: P2 (perinatal), P10 (neonatal), P21 (weaning), P42 (sexual maturity), and P84 (adult) from proximal pulmonary artery-derived cells from male C57BL/6J mice, yielding 11,143 cells in total (**Fig. 1A**). Uniform Manifold Approximation and Projection (UMAP) revealed distinct trajectories of major vascular cell populations including endothelial cells (ECs), smooth muscle cells (SMCs), fibroblasts (FBs), and Macs, annotated using consensus markers from CellMarker2, PanglaoDB, and CellXGene **(Fig. 1B**). Major vascular cell populations—including endothelial cells (ECs), smooth muscle cells (SMCs), fibroblasts (FBs), and Macs—were annotated using consensus markers from CellMarker2, PanglaoDB, and CellXGene (**Fig. 1C** and **Extended Data Fig. 1**). Cell population dynamics from P2 to P84 confirmed marked shifts up to sexual maturity (P2–P42), followed by stabilization consistent with the establishment of adult vascular homeostasis (**Fig. 1D**). These findings align with prior evidence that biomechanical maturity emerges in proximal pulmonary arteries around P34^21^.

**Figure 1.**
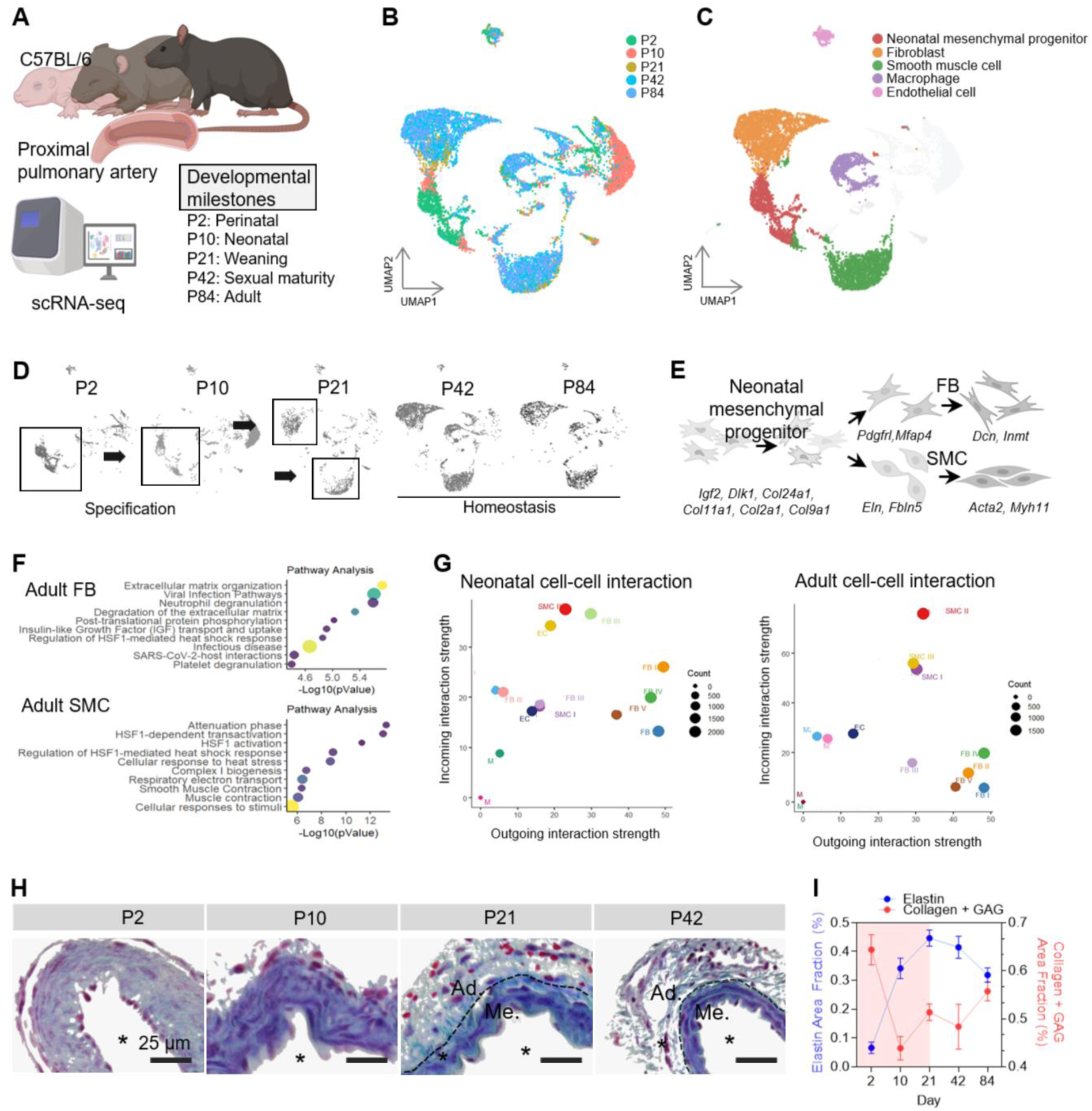
Transcriptomic and cellular phenotype changes in major vascular cell types of the developing mouse pulmonary artery closely parallel tissue organization and remodeling events. (A) Schematic representation of mouse pulmonary artery development from the neonatal stage (P2) to adulthood (P84), illustrating key developmental milestones. UMAP visualization of single-cell RNA sequencing data from pulmonary artery cells at (B) all developmental ages (3-5 mice each), depicting (C) five major cell populations, including EC, FB, SMC, Mac, and a neonatal mesenchymal progenitor population. (D) UMAP visualization depicting cellular changes with age (P2, P10, P21, P42, and P84). Black boxes highlight regions indicating dynamic changes in cellular composition. (E) Schematic model illustrating the specification of the neonatal progenitor populations into FBs and SMCs. Gene names represent age- and cluster-specific markers associated with respective differentiation pathways. (F) Pathway enrichment analysis comparing neonatal and adult subpopulations for each lineage: FB and SMC. Enrichment was performed using Reactome databases on differentially expressed genes. Dot size indicates gene ratio; color corresponds to significance (–log₁₀(p- value)). (G) Cell-cell interaction pattern in neonatal and adult stages, showing shifts in sender-receiver patterns. The x-axis represents outgoing interaction strength, and the y-axis represents incoming interaction strength. Bubble size indicates interaction count. Data are presented as mean ± SEM. (H) Masson’s Trichrome–stained histology images showing pulmonary artery structural remodeling from P2 to P42. The vessel transitions from a double-layered structure (P10), with EC and FB-SMC progenitor, to a three-layered configuration (P21,P42), with EC, FB, and SMC contributing to vascular wall formation. *: Lumen, Me.: Tunica media, Ad.: Tunica adventitia. Scale bar = 25 μm. (I) Quantification of elastin, collagen and cytoplasm area fractions across developmental stages, demonstrating shifts in extracellular matrix composition (n = 4-6).

A transient FB–SMC–like population enriched at P2 was particularly noteworthy (boxed region, **Fig. 1D**). By P21, this cluster resolved toward annotated fibroblast and smooth muscle cell lineages (**Fig. 1D, E**), consistent with a transitional state unique to early postnatal pulmonary artery development. Crnkovic *et al.* reported that FB and SMC lineages are transcriptomically linked and can interconvert in adult artery remodeling contexts^24^, but the emergence of a defined FB–SMC population specifically during the neonatal period represents a previously unrecognized transitional state enabled by time-course single-cell analysis. This cluster co-expressed canonical FB and SMC markers (*Dcn* and *Acta2*, respectively^24, 25^) (**Extended Data Fig. 2A**) and was enriched for progenitor-associated genes, including *Dlk1*, a known marker of immature mesenchymal progenitors in multiple tissues^26^, and *Igf2* as well as cartilage-associated collagens (*Col11a1*, *Col2a1*, *Col24a1*). These cells transitioned toward lineage-restricted fates marked by downregulation of early ECM-remodeling genes (*Col11a1*, *Pdgfrl*, *Col24a1*, *Fbln5*, *Eln*) and induction of mature FB (*Mfap4*, *Dcn*, *Inmt*) or SMC markers (*Acta2*, *Myh11*) (**Fig. 1E**, **Extended Data Fig. 2B**, C)^25, 27–29^. This specification parallels evidence that pericardial mesenchyme can give rise to both SMCs and FBs in the coronary artery, now extended to the proximal pulmonary artery^30^. By directly sampling neonatal stages (P2–P10), this dataset provides time-resolved transcriptomic evidence for a transient FB–SMC population unique tot he neonatal pulmonary artery, offering new insight into vascular lineage specification during postnatal arterial growth. To our knowledge, this is the first time-course single-cell transcriptomic analysis across defined postnatal windows to elucidate proximal pulmonary artery development.

We next examined transcriptional maturation across the major vascular cell types. The FBs were enriched for ECM biosynthesis programs up to P21, whereas adult FBs transitioned to matrix remodeling (e.g., *Mmp2*, *Col8a1*), stress-response (*Hsf1*), and inflammatory signaling modules (**Fig. 1F**, **Extended Data Fig. 3A**, B). Concurrently, the SMCs shifted from more of a synthetic to a contractile phenotype, characterized by upregulation of mitochondrial and oxidative phosphorylation genes (**Fig. 1F**, **Extended Data Fig. 3A**, C). The Macs transitioned from canonical pro-inflammatory states (*Lyz2*, *S100a9*, *Ccl8*) to an anti-inflammatory regulatory phenotype characterized by expression of *Il10*, MHC-II, and cholesterol-handling genes (**Extended Data Fig. 3A**, D).

To understand how transcriptional maturation reshaped cell–cell communication, we analyzed receptor-ligand interactions using CellChat (v2)^31^. At P2, during the early postnatal period, cells exhibited high signaling diversity with broad cross-talk among mesenchymal and immune populations. By adulthood at P84, cell–cell signaling became more specialized and modular, predominantly featuring selective pathways (e.g., IL6, IGFBP, and CLEC7A), likely reflecting functional specialization of mature vascular compartments (**Extended Data Fig. 4**). This shift was accompanied by emerging role specialization—FBs transitioned into primary signaling hubs, whereas SMCs predominantly received cues (**Fig. 1G**). These findings suggest that postnatal development not only alters cellular states, it also reorganizes the communication network itself to enable a division of labor and modular control in maturity.

This cellular specification coincided with a structural transition into a distinctive three-layered wall. Histological analysis revealed coordinated changes in pulmonary artery architecture during postnatal development (**Fig. 1H**). At P2, the vessel consisted primarily of a mixture of cells and fibrillar collagen with no clear medial lamellar structure. By P10, the vessel included an endothelial layer plus a loosely organized, elastin-rich medial zone with sparse adventitial collagen. By P21, all three layers – intima, media, and adventitia – were distinct histologically, with the medial and adventitial layers each populated by a single dominant cell type, SMCs and FBs, respectively. The medial layer became more condensed and circumferentially aligned between P21 and P42 while the adventitia expanded, consistent with increased structural support despite the lack of increase in pulmonary artery pressure (**Extended Data Fig. 5A**). Alongside the structural changes, there was a dynamic remodeling in ECM composition. The elastin fraction increased, peaking at P21 before declining thereafter. By contrast, collagen and glycosaminoglycan (GAG) content showed an inverse trend relative to elastin (**Fig. 1I**), consistent with maturation of an elastin-driven elasticity with collagen-mediated reinforcement. In parallel, the cellular area fraction dropped significantly from P2 to P21 (*P* < 0.001), with associated medial compaction and growth of the collagen-rich adventitia (**Extended Data Fig. 5B**). Together, these data identify P21 as a morpho-functional inflection point for arterial maturation, marked by dramatic changes in cell phenotype and ECM architecture.

### CausaLink integrates unpaired transcriptomics and morpho-mechanical data

To resolve causal relationships without paired samples, we developed CausaLink, a framework that integrates unpaired transcriptomic and biomechanical data. CausaLink operates through three key modules: (1) a cross-modal autoencoder that learns a shared latent representation between gene expression and biomechanical traits without requiring one-to-one sample pairing; (2) a cross-scale score-based causal discovery algorithm that infers directed acyclic graphs (DAGs) linking prioritized genes to vascular properties; and (3) *in silico* simulations that predict system-level remodeling outcomes following gene perturbation (**Fig. 2A**).

**Figure 2.**
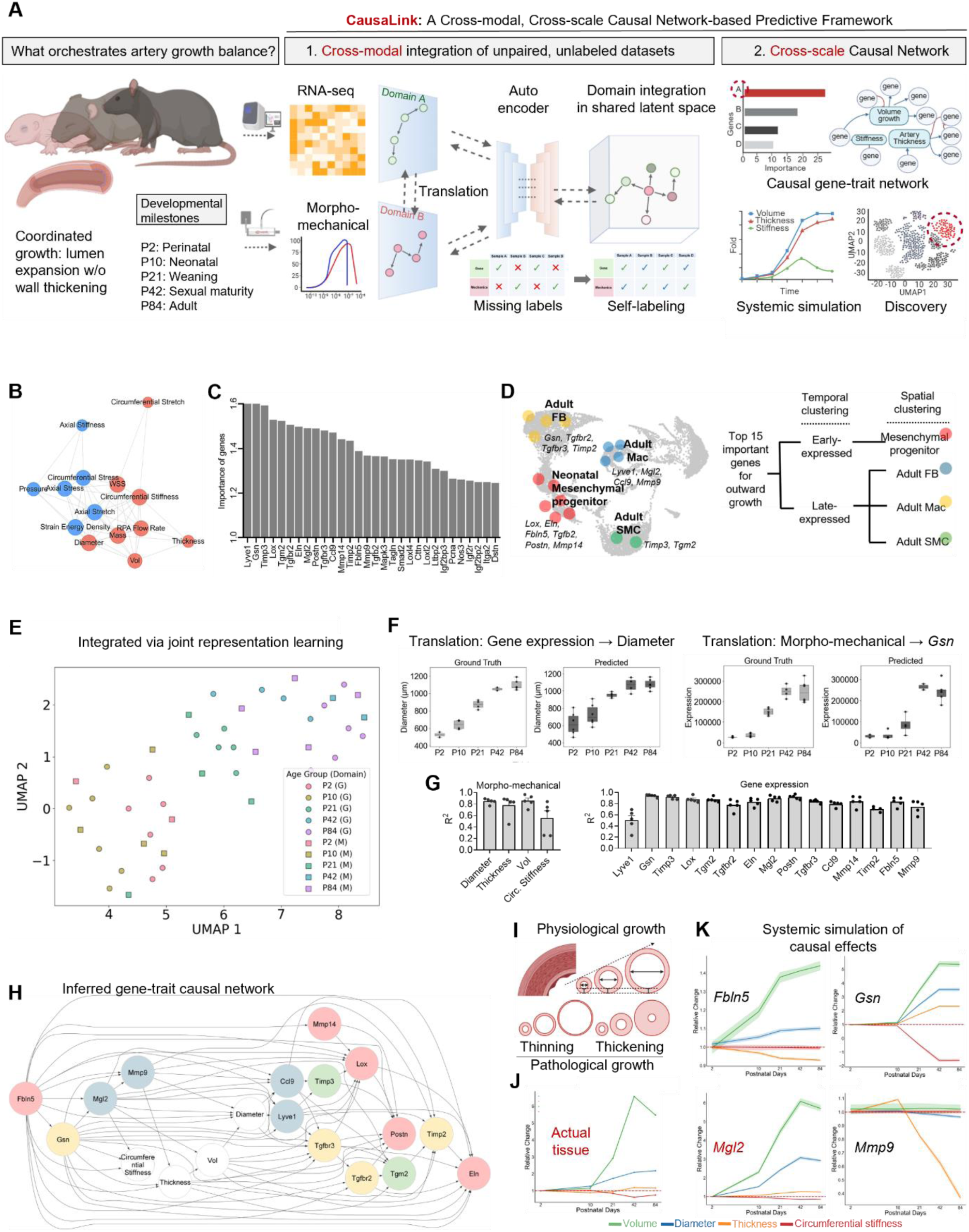
CausaLink pinpoints *Mgl2*+ Macs as a main orchestrator for balanced artery growth by integrating unpaired and unlabeled datasets to infer causal gene-trait network. (**A**) Schematic illustration to describe the causal integration framework. (**B**) Network analysis illustrating relationships among mechanical parameters in the arterial wall. Two clusters are highlighted in red and blue. (**C**) Top 30 important genes contributing to circumferential stiffness change, ranked based on feature importance scores from PLSR. (**D**) Mapping and categorizing the top 15 genes correlated with circumferential stiffness across neonatal mesenchymal progenitor, adult FB, adult SMC, and adult Mac populations, identifying key cell types involved in ECM remodeling. (**E**) Shared latent space that integrate transcriptomic and tissue properties domain via joint representation learning. G; Transcriptomic data, M: mechanical and morphological data. (**F**) Bidirectional translation between transcriptomic and morpho- mechanical domains. Left, ground-truth versus predicted arterial diameter across postnatal stages (P2–P84) from gene expression input. Right, ground-truth versus predicted expression of *Gsn* across postnatal stages from biomechanical input. Predicted trajectories closely recapitulated temporal trends observed in the original datasets. (**G**) Quantitative assessment of translation fidelity. Bar plots show coefficient of determination (R²) values between predicted and ground-truth measurements for morpho- mechanical traits (diameter, thickness, volume, and circumferential stiffness) and selected vascular remodeling genes (*Lyve1, Gsn, Timp3, Lox, Tgm2, Tgfbr2, Eln, Mgl2, Postn, Tgfbr3, Ccl9, Mmp14, Timp2, Fbln5,* and *Mmp9*). Data are presented as individual samples with mean ± SEM. N = 5. (**H**) DAG illustrating cause-and-effect relationships between the top 15 key genes (colored by cell type) and mechanical properties involved in pulmonary artery outward growth (white). Edge arrows indicate causal directionality from cause to effect. Nodes are grouped by cell type origin, including neonatal mesenchymal progenitor (yellow), FBs (Green), SMCs (Red), and Macs (Blue). (**I**) Schematic illustration of normal and pathological artery growth patterns, depicting balanced growth in normal conditions and wall-thinning or wall-thickening in pathological conditions. (**J**) Measured morpho- mechanical property changes across postnatal development, showing relative changes in diameter, thickness, volume, and circumferential stiffness of the pulmonary artery. (**K**) Causal effect simulation of four key driver genes (*Fbln5*, *Gsn*, *Mgl2*, and *Mmp9*) on tissue properties, illustrating their predicted influence on diameter, thickness, and circumferential stiffness across postnatal development.

We first prioritized biologically relevant features by identifying genes and tissue traits associated with vascular development. From 14 morphometric and mechanical parameters quantified in the right pulmonary artery of 27 mice (4-6 mice per stage), network analysis revealed two major trait clusters. One cluster mapped to progressive luminal enlargement, with circumferential stiffness emerging as a central hub (**Fig. 2B**), consistent with the concept that appropriate circumferential stiffness permits radial growth during arterial development. To connect genes with trait, partial least squares regression (PLSR) integrating RNA-seq with measured circumferential stiffness prioritized 15 genes with the highest variable importance: *Lyve1, Gsn, Timp3, Lox, Tgm2, Tgfbr2, Eln, Mgl2, Postn, Tgfbr3, Ccl9, Mmp14, Timp2, Fbln5,* and *Mmp9* (**Fig. 2C**). Their diverse temporal expression across neonatal mesenchymal progenitors, FBs, SMCs, and Macs, indicates a distributed, lineage-spanning regulatory network (**Fig. 2D**).

As a first step, we integrated unpaired gene expression and tissue trait measurements using a cross-modal autoencoder architecture comprising dual encoders and decoders. Each encoder independently projects transcriptomic or biomechanical data into a shared latent space, and subsequently, the decoders reconstruct each original modality from this shared representation. Two additional losses optimize the model: cycle-consistency enforces preservation of representations during cross-modal translation and back-translation, while maximum mean discrepancy (MMD) aligns the latent distributions to minimize modality-specific bias. To evaluate reconstruction fidelity, we compared original and reconstructed data in both domains. PCA revealed that reconstructed gene expression profiles closely matched the original variance structure (**Extended Data Fig. 6A**), while biomechanical traits—including diameter, thickness, mass, volume, and circumferential stiffness—were accurately reconstructed across samples (**Extended Data Fig. 6B**). These results demonstrated that the latent space preserves essential biological information from each modality, providing a robust foundation for downstream causal inference. Importantly, joint representation learning aligned transcriptomic and biomechanical data by developmental stage rather than modality, with UMAP embedding confirmed coherent cross-modality co-clustering across ages (**Fig. 2E**). This robust integration indicates that the model captures shared developmental programs across scales, enabling cross-domain causal analysis.

This architecture supports bidirectional translation, whereby gene expression profiles predicted vascular biomechanical traits and, conversely, biomechanical features reconstructed gene expression trajectories (**Fig. 2F**). Cross-domain predictions achieved high fidelity (R² = 0.7–0.9 for diameter, thickness, and volume) and moderate accuracy for circumferential stiffness (R² ≈ 0.5). Reconstructed trajectories of key vascular remodeling genes—including *Lyve1*, *Lox*, *Eln*, *Mgl2*, *Postn*, and *Mmp9*—also reached R² values of 0.7–0.9 (**Fig. 2G**), reflecting consistent recovery of ECM- and immune-related programs. These results demonstrate that the shared latent space supports accurate, quantitative cross-modal inference, extending beyond qualitative trend preservation to predictive modeling of biologically meaningful genes and traits. Beyond alignment, this translation module in CausaLink offers a framework to infer transcriptomic profiles from abundant or retrospectively pooled phenotypic datasets—a capability increasingly valuable in multimodal deep learning^32^.

### CausaLink pinpointed *Mgl2*^+^ Macs as an orchestrator of balanced growth

We next applied a score-based causal discovery using the Discovery At Scale (DAS) algorithm to infer a directional gene–trait network. DAS is particularly suitable for biological datasets, as it robustly captures nonlinear relationships and remains reliable despite violations of conventional statistical assumptions^33, 34^. Using the integrated CausaLink dataset, we constructed a DAG linking 15 prioritized genes to four key morpho-mechanical features: luminal diameter, wall thickness, circumferential stiffness, and wall volume as an integrated metric of overall growth (**Fig. 2H**). The resulting DAG identified four upstream regulators with multi-trait influence, including genes encoding elastin-associated fibulin-5 (*Fbln5*), actin-binding protein gelsolin (*Gsn*), matrix metalloproteinase 9 (*Mmp9*), and macrophage galactose N-acetyl-galactosamine specific lectin 2 (*Mgl2*). The fact that only 4 out of 15 correlated genes were inferred as primary upstream drivers underscores the limitation of correlation-based models and conventional machine learning approaches, reinforcing the necessity of causal inference to uncover true mechanistic regulators.

Having established the causal hierarchy of key genes, we next evaluated whether the inferred relationships could quantitatively reproduce known physiological remodeling outcomes. System-wide simulations of gene perturbations yielded predictions that highly consistent with known biological outcomes. For example, simulated suppression of *Fbln5*— which mirrors its progressive downregulation during postnatal development—was predicted to enhance luminal diameter and wall volume expansion, albeit to a lesser extent than observed *in vivo*^35–37^. This prediction aligns with a known role of *Fbln5* in elastic fiber assembly, as *Fbln5*- deficient mice exhibit compromised elastic fiber organization, vessel tortuosity, and dilation, leading to vascular dysfunction, supporting the predictive validity of the causal model. Simulated upregulation of *Mmp9* induced wall thinning, consistent with its known function in ECM degradation and aneurysm formation^18^. These congruencies support the physiological relevance of the inferred causal framework.

Notably, *Mgl2*, orthologous to human CLEC10A, was inferred as a regulator of balanced arterial growth. Simulated 7.1-fold increases in *Mgl2* expression led to proportional enlargement of vessel diameter (∼3-fold) and wall volume (∼5.8-fold) without altering arterial wall thickness and stiffness, closely matching experimentally observed changes *in vivo* (2.2- fold and 5.5-fold at P84, respectively) (**Fig. 2J, K**). While *Mgl2* has been widely studied in immunoregulatory contexts^38–41^, its role in orchestrating vascular structure during normal development remains poorly characterized. These findings uncover a previously unrecognized developmental function of *Mgl2*⁺ Macs in promoting homeostatic arterial growth and remodeling.

To further investigate this regulatory mechanism, we examined *Mgl2* expression dynamics during postnatal development. Although per-cell *Mgl2* expression levels remained relatively stable, the proportion of *Mgl2*⁺ cells increased substantially from P2 to P84, resulting in a net rise in tissue-level *Mgl2* expression over time (**Fig. 3A-C**). Co-expression analysis with the immune cell marker *Ptprc* confirmed a selective expansion of the Mgl2⁺ Mac subpopulation (**Fig. 3D**). *Mgl2*⁺ Mac in our dataset comprise both *Lyve1*⁺ and *Ccr2*⁺ subsets, corresponding to tissue-resident homeostatic and monocyte-derived Macs, respectively^42, 43^ (**Fig. 3E, F**). Notably, the *Lyve1*⁺ subset constitutes the majority (0.5–0.8), whereas *Ccr2*⁺ Macs represented a smaller fraction (0.07–0.18). Across these populations, both anti-inflammatory (*Mrc1*, *Il10*, *Cd163*, and *Ccl24*) and pro-inflammatory genes (*Tnf*, *Il1b*, *Il6*, and *Cd80*) were expressed (**Fig. 3G, H**), consistent with the established roles of *Lyve1*⁺ and *Ccr2*⁺ lineages in previous studies^42, 44, 45^. Importantly, from P10 to P21, the proportions of *Lyve1*⁺ and *Ccr2*⁺ cells within the *Mgl2*⁺ compartment fluctuated dynamically, before stabilizing by P42–P84—a temporal pattern that corresponds with periods of vessel enlargement. Together, these results reveal dynamic shifts in the proportions of *Lyve1*⁺ and *Ccr2*⁺ subtypes within the *Mgl2*⁺ Mac population during vessel growth, providing insight into how compositional balance among macrophage lineages may contribute to coordinated tissue remodeling alongside the established homeostatic roles for Lyve1⁺ macrophages^46, 47^.

**Figure 3.**
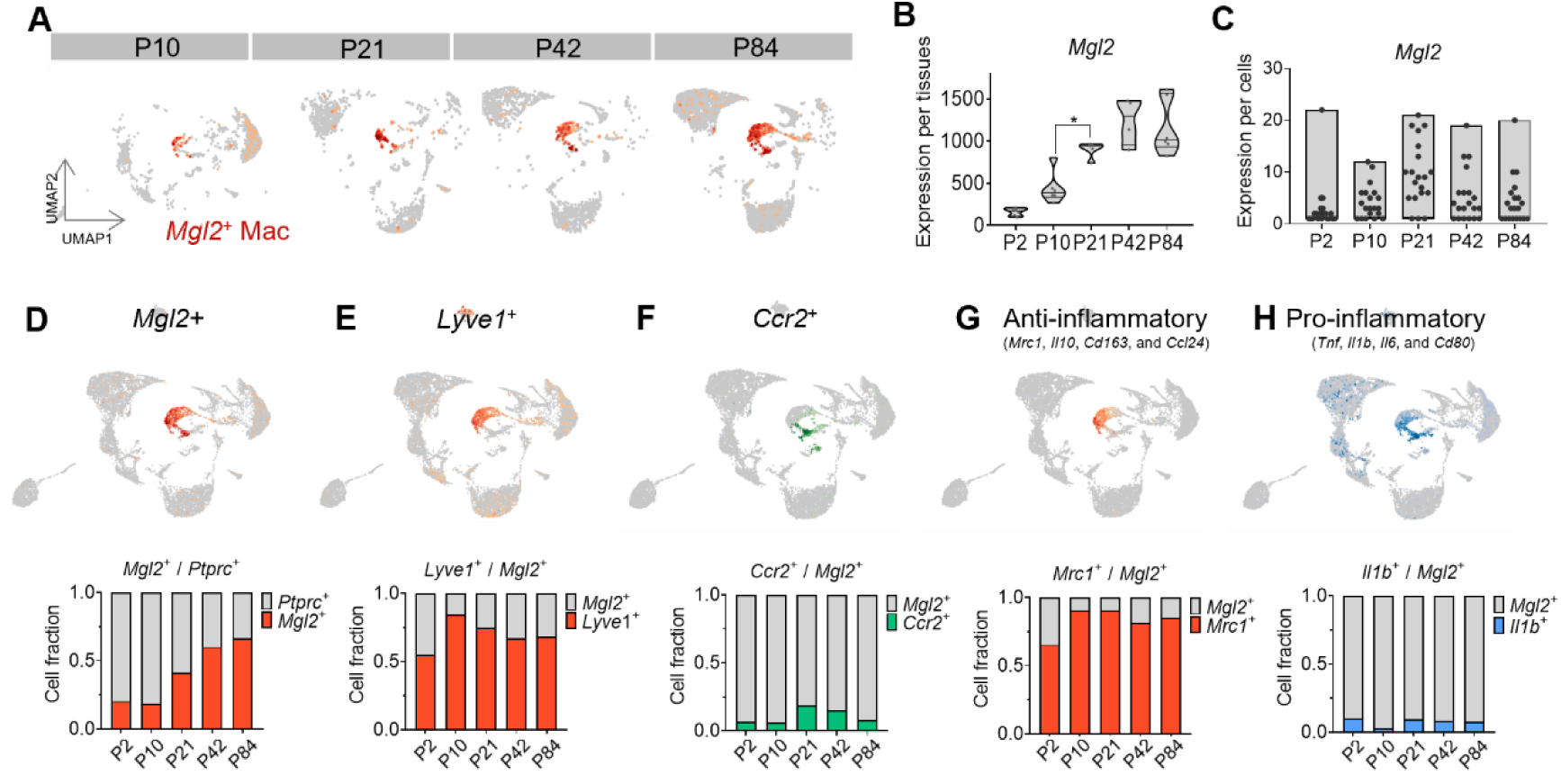
Time-course single-cell mapping of *Mgl*2⁺ macrophage dynamics and polarization during postnatal pulmonary artery growth. (**A**) Feature plots of *Mgl2+* cells (red) across postnatal times (P2–P84), showing expression intensity and spatial distribution during postnatal development. Color represents expression intensity. *Mgl2* gene expression (**B**) per tissue and (**C**) per cell across developmental stages. Data are presented as mean ± SEM. A one-way analysis of variance and Tukey’s test were performed for statistical analysis (\**P* < 0.05). (**D**) Changes in the proportion of *Mgl2+* Macs among *Ptprc^+^* immune cells across five developmental stages, illustrating macrophage proliferation during postnatal artery growth. Feature plots of (**E**) *Lyve1^+^*, (**F**) *Ccr2^+^*, (**G**) Anti-inflammatory and, and (**E**) pro-inflammatory marker expressing macrophages. Stacked bar charts depict the fraction of *Mgl2*⁺ macrophages co-expressing the indicated markers across developmental stages.

### MGL^high^ macrophages orchestrate balanced vascular remodeling in human arterial assembloids

Guided by CausaLink predictions that *Mgl2*⁺ Macs coordinate arterial remodeling by expanding lumen while preserving wall thickness, (**Fig. 2**) we co-cultured hiPSC-derived blood vessel organoids (hBVOs) with THP-1-derived Macs polarized into pro-inflammatory MGL^low^ or anti- inflammatory MGL^high^ states (**Fig. 4A**). Macs were introduced during the mural-cell differentiation phase to restore immune cues absent in standard *in vitro* organoids. hBVOs recapitulate endothelial-mural organization^48^ and yield quantifiable readouts of lumen diameter and mural thickness^49, 50^, precisely the trait pair that operationalizes balanced remodeling in our framework, thereby providing a physiologically relevant human context for experimental validation. Recent scRNA-seq analyses revealed significant differences between *in vitro* and transplanted hBVOs^51^, namely, increased immune-related signaling and decreased TGF-β signaling activity in transplanted organoids, suggesting that additional immune cues are required for proper vascular specification.

**Figure 4.**
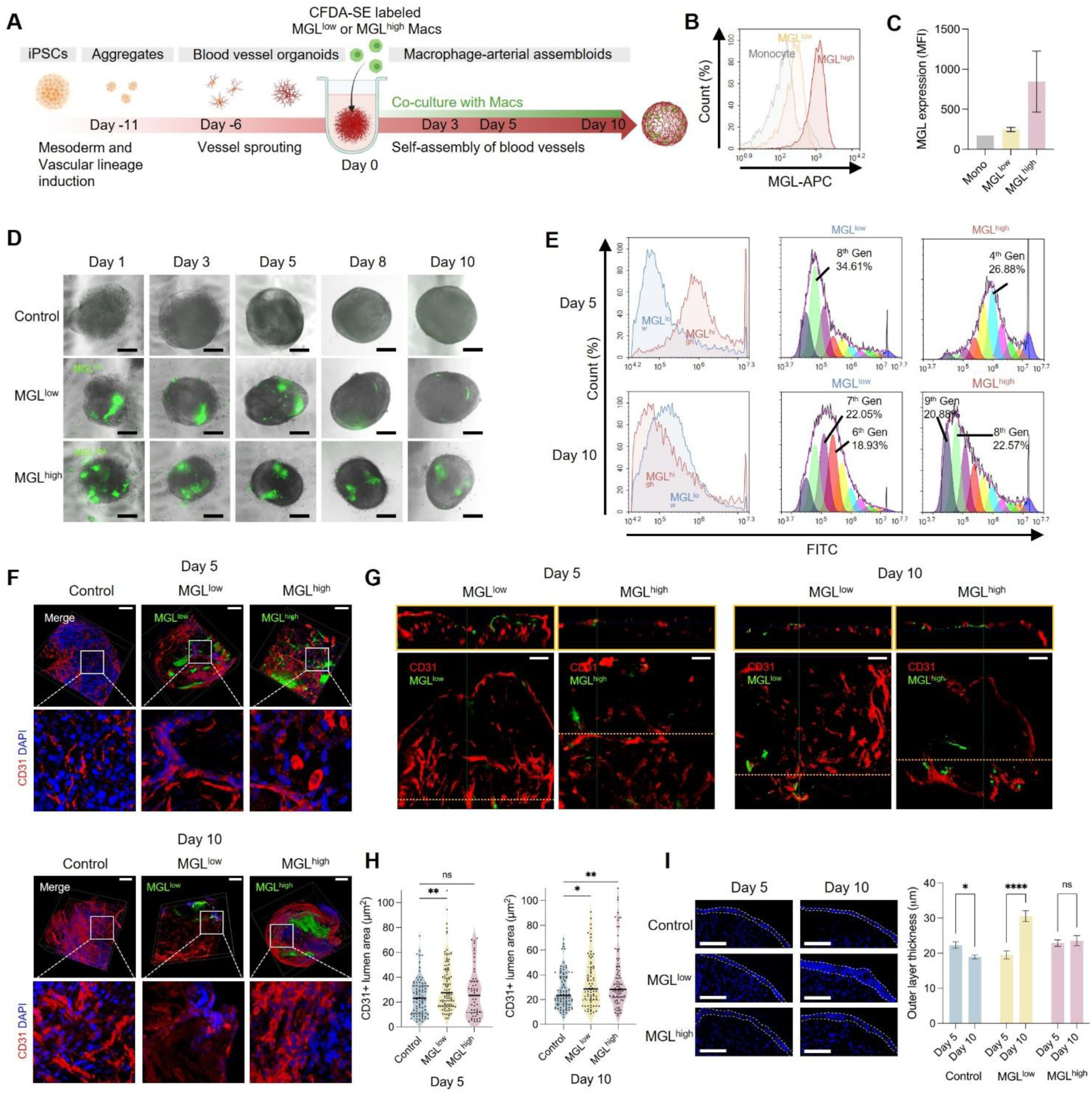
MGL^high^ Macs orchestrated balanced growth in developing arterial assembloids. (A) Schematic illustration of generation of macrophage (Mac)-arterial assembloids. Co-cultured Macs migrated into blood vessel organoids and influenced vascular remodeling during the self-assembly of blood vessels. (B) Overlay histograms showing MGL expression in THP-1 monocytes (Monocyte), THP-1- derived MGL^low^ Macs (MGL^low^), and THP-1-derived MGL^high^ Macs (MGL^high^). (C) Mean fluroscence intensity of MGL in THP-1 monocytes, MGL^low^, and MGL^high^ Macs analyzed by flow cytometry. (D) Live images of assembloids co-cultured with CFDA-SE-labeled Macs from Day 1 to Day 10. Scale bars = 200 μm. (E) Left; overlay histograms of CFDA-SE fluorescence in MGL^low^ and MGL^high^ Macs in assembloids at Day 5 and Day 10. Right; analysis showing rapid proliferation of MGL^low^ Macs compared to MGL^high^ on Day 5, followed by sustained proliferation of MGL^high^ Macs on Day 10. The most abundant cell generation is indicated at the peak. (F) Reconstructed three-dimensional CD31^+^ microvessel structures in assembloids at Day 5 and Day 10. Zoomed-in images highlight morphological differences of ECs across experimental groups. Scale bars = 50 μm. (G) Orthogonal views showing co-cultured Macs and CD31^+^ microvessels in assembloids at Day 5 and Day 10. (H) Comparison of lumen areas in assembloids co-cultured with MGL^low^ or MGL^high^ Macs versus control, showing significant Mac-driven increases in vessel diameter. Data are presented as mean ± SEM. Statistical significance was determined by one-way ANOVA with Dunnett’s post hoc test for multiple comparisons against the control. (I) Changes in outer layer thickness between Day 5 and Day 10 assembloids, demonstrating abnormal arterial growth in the absence of MGL^high^ Macs. Data are presented as mean ± SEM. Statistical significance was determined by two-way ANOVA with Sidak’s post hoc test. ****p < 0.0001; ***p < 0.001; **p < 0.01; *p < 0.05; ns, not significant.

To address this signaling gap, we incorporated MGL^high^ and MGL^low^ Mac subsets into the assembloids during the mural cell differentiation. We selected THP-1-derived Macs to ensure scalability and batch-to-batch reproducibility required for cross-scale validation. Although THP-1 cells and primary human Macs differ in ontogeny and polarization capacity, THP-1-derived Macs provide standardized differentiation, a stable genetic background, and minimal donor variability^52^. We therefore used these cells to interrogate the directionality of Mac effects with higher reproducibility, while interpreting effect sizes in light of these lineage differences.

To establish MGL-dependent Mac subsets, THP-1 monocytes-derived M0 Macs were polarized into pro-inflammatory (MGL^low^) and anti-inflammatory (MGL^high^) states using interferon-γ (IFN-γ) with lipopolysaccharide (LPS) or IL-10 with TGF-β1, respectively. These subsets exhibited low and high expression of MGL genes and proteins, as confirmed by qRT-PCR and flow cytometric analyses (**Figs. 4B, C**). This design enabled a direct comparison of MGL-dependent effects on vascular growth in a human cell-based 3D vascular model. During *in vitro* differentiation, hBVOs formed microvessels consisting of endothelial lumen surrounded by a mural layer, allowing real-time observation of immune–vascular crosstalk during remodeling. The introduced Macs actively migrated toward hBVO-derived vessels, influencing endothelial and mural differentiation during the subsequent self-assembly phase and inducing structural differences in the assembloids.

We first examined the dynamics of co-cultured Macs in assembloids as a function of MGL expression level. Live-cell imaging over 10 days revealed distinct proliferative dynamics between Mac subsets (**Fig. 4D**). Specifically, MGL^low^ Macs proliferated rapidly, undergoing up to eight cell divisions by Day 5, but subsequently ceased proliferation and showed subsequent decline in abundance (**Fig. 4E)**. In contrast, MGL^high^ Macs showed slower yet sustained proliferation throughout 10 days, consistent with a stable, homeostatic role in tissue remodeling, in contrast to the transient inflammation-associated expansion of MGL^low^ subsets. These findings support our murine scRNA-seq data in developing tissues, revealing a neonatal predominance of *Mgl2^-^* immune cells followed by rapid expansion and eventual dominance of *Mgl2^+^* subsets during arterial maturation (**Fig. 3D**).

Next, we examined structural changes in the assembloids according to MGL expression. Immunofluorescence imaging revealed significant differences in endothelial network architecture and microvessel organization between assembloids containing MGL^low^ and MGL^high^ Macs (**Fig. 4F, G**). By Day 5, assembloids co-cultured with MGL^low^ Macs exhibited extensive but disorganized angiogenesis, characterized by irregular vessel branching indicative of pathological remodeling. By Day 10, decline in MGL^low^ Macs coincided with abnormal microvessel overgrowth. In contrast, assembloids containing MGL^high^ Macs initially showed limited EC alignment but developed robust, well-organized microvascular networks by Day 10, consistent with predictions from our causal modeling. The close association of MGL^high^ Macs with microvessels suggests a direct role in promoting stable and controlled angiogenesis. Orthogonal projections confirmed that both MGL^low^ and MGL^high^ Macs localized in close proximity to CD31^+^ microvessels (**Fig. 4G**). Morphometric analysis further revealed subset- specific effects on vessel lumen growth (**Fig. 4H**). By Day 10, assembloids with MGL^high^ Macs demonstrated significantly increased vessel diameters and improved vascular organization, reflecting balanced, physiologically relevant remodeling compared to controls and MGL^low^ Mac assembloids.

Additionally, mural outer layer thickness remained stable only in MGL^high^ Mac assembloids, further indicating balanced vascular remodeling. We next assessed cellular dynamics within the outer mural region resembling a vascular wall of the assembloids, which includes outer mural layer resembling a vascular wall (**Fig. 4I**). In line with previous observations, control assembloids showed reduced mural thickness from Day 5 to Day 10, whereas assembloids co-cultured with MGL^low^ Macs showed marked mural thickening, indicative of pathological remodeling. In contrast, mural thickness remained stable in MGL^high^ Mac assembloids despite increased vessel diameters, suggesting balanced arterial remodeling.

### Macrophage-mediated mural lineage diversification and ECM remodeling

Next, we assessed Mac subset-specific roles in mural cell differentiation and ECM remodeling using the assembloid model. Initially, assembloids self-organized into three- dimensional spherical structures, comprising distinct outer endothelial and inner mesenchymal layers containing SMCs and FBs^53^ (**Fig. 5A**). The cellular architecture of this outer region changed dynamically during the self-assembly process, with CD31^+^ ECs forming the outermost luminal lining, while PDGFRβ^+^ mesenchymal cells assembled into subjacent sub-endothelial layers (**Extended Data Fig. 9**). Assembloids containing MGL^low^ Macs predominantly exhibited expansion of αSMA^+^ SMC layers, whereas those with MGL^high^ Macs initially promoted the formation of PDGFRα^+^ FB layers and subsequently maintained stable thickness of both SMC and FB layers through Day 10 (**Fig. 5 B-E, Extended Data Fig. 10A**). Analysis of the relative fractions of SMC and FB layer thicknesses within the outer mesenchymal layer further supported these observations (**Fig 5F**). This remodeling pattern closely paralleled the maturation observed in mouse pulmonary arteries from P21 to P42 (**Extended Data Fig. 5A**), suggesting a conserved mechanism driven by MGL^high^ Macs. The FB fraction (∼70%) in MGL^high^ Mac assembloids was highly consistent with the adventitial fraction measured *in vivo* after P21 (**Extended Data Fig. 5A**), further supporting a homeostatic role for MGL^high^ Macs in vascular development.

**Figure 5.**
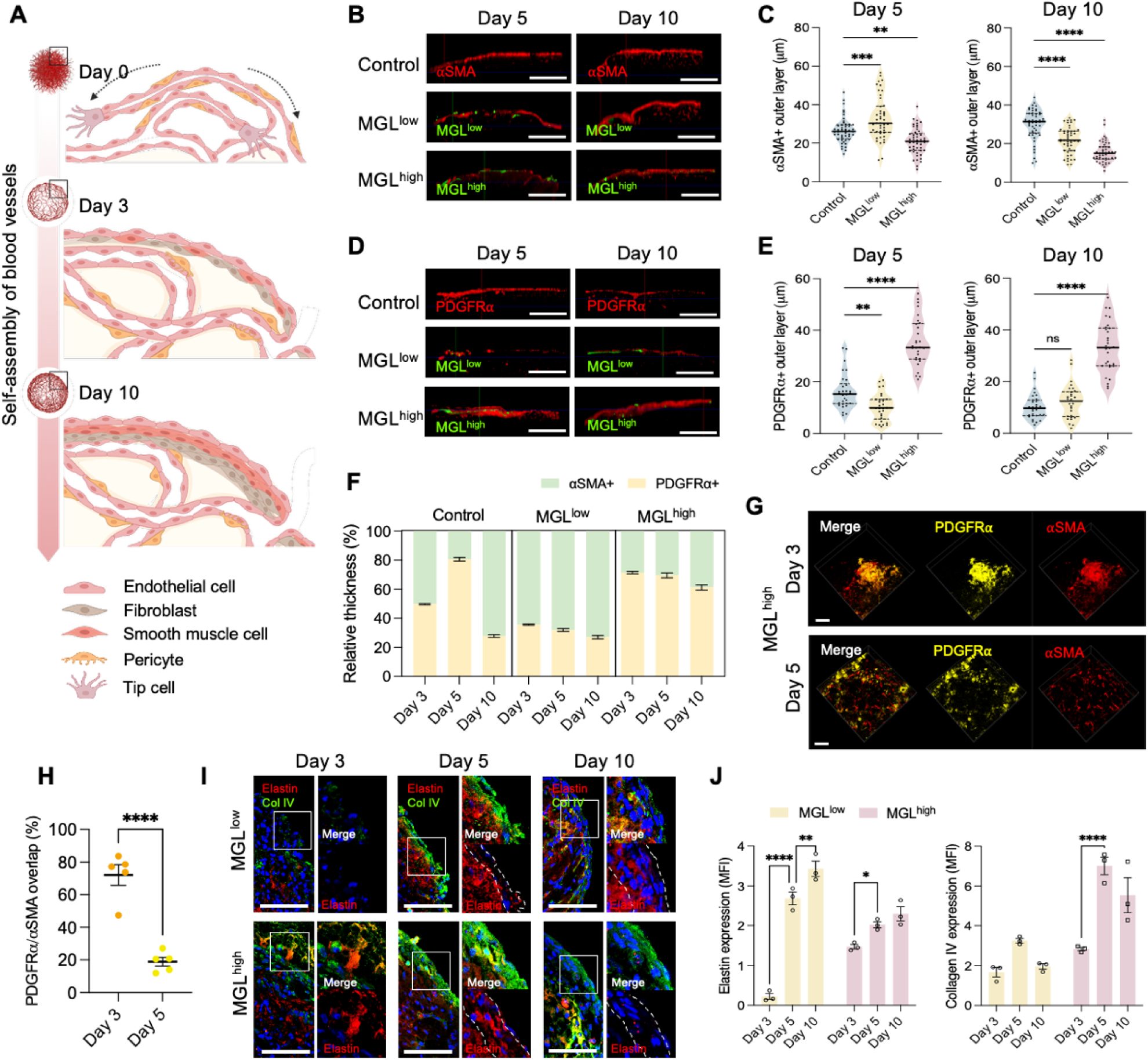
MGL^high^ Macs promote neonatal mesenchymal progenitor and FB development to maintain mesenchymal layer thickness in arterial assembloids. (A) Schematic illustration of the self-assembly of blood vessels in the outer region of macrophage (Mac)-arterial assembloids. The outer layer consists of an outermost endothelial layer and a mesenchymal layer, which includes both smooth muscle cells (SMCs) and fibroblasts (FBs). (B) Orthogonal views showing CFDA-SE-labeled Macs and outer αSMA^+^ SMC layers in assembloids at Day 5 and Day 10. Scale bars = 200 μm. (C) Comparison of αSMA^+^ layer thickness in assembloids co-cultured with MGL^low^ or MGL^high^ Macs versus control, showing significant MGL^high^ Mac-driven increases in SMC thickness at Day 10. Data are presented as mean ± SEM. Statistical significance was determined by one-way ANOVA with Dunnett’s post hoc test for multiple comparisons against the control. (D) Orthogonal views showing CFDA-SE-labeled Macs and PDGFRα^+^ FB layers in assembloids at Day 5 and Day 10. Scale bars = 200 μm. (E) Comparison of PDGFRα^+^ layer thickness in assembloids co-cultured with MGL^low^ or MGL^high^ Macs versus control, showing significant MGL^low^ Mac-driven increases in FB thickness at Day 5. Data are presented as mean ± SEM. Statistical significance was determined by one-way ANOVA with Dunnett’s post hoc test for multiple comparisons against the control. (F) Changes in the relative proportion of outer mesenchymal layer thickness over time during Mac co-culture, illustrating distinct patterns of arterial growth driven by MGL^ow^ or MGL^high^ Macs. (G) Populations co-expressing PDGFRα and αSMA were observed in assembloids co-cultured with MGL^low^ Mac at Day 3. (H) The overlap between PDGFRα and αSMA expression significantly decreased by Day 5. (I) Time-dependent changes by MGL^low^ or MGL^high^ Macs in elastin and collagen IV expression in the outer region. Scale bars = 50 μm. (J) Changes in elastin and collagen type IV expression in assembloids during Mac co-culture, in line with MGL^low^ or MGL^high^ Mac-induced remodeling of mesenchymal layer composition. Data are presented as mean ± SEM. Statistical significance was determined by two-way ANOVA with Sidak’s post hoc test. ****p < 0.0001; ***p < 0.001; **p < 0.01; *p < 0.05; ns, not significant.

We further validated Mac-driven mural lineage specification, a critical developmental milestone predicted by scRNA-seq analyses (**Fig. 1D, E**). At Day 3, PDGFRα^+^ FB and αSMA^+^ SMC co-localized within MGL^high^ assembloids, consistent with a neonatal mesenchymal progenitor state observed *in vivo*. Distinct spatial segregation of these lineages was evident by Day 5, with PDGFRα^+^ FBs predominantly localized to the outer adventitial-like region and αSMA^+^ SMC layers aligning closer to the endothelial layer, mimicking native arterial wall organization and confirming Macs involvement in lineage specification during arterial maturation (**Fig. 5G-H, Extended Data Fig. 10B**). Notably, co-culture with MGL^high^ Macs induced significant adventitial FB expansion, a hallmark of later-stage arterial maturation, mirroring *in vivo* developmental dynamics. Consistent with previous reports showing increased mural FB populations in transplanted hBVOs^51^, these findings further support the role of Mac- enriched environments in FB expansion and maturation. Importantly, our assembloid model overcomes critical limitations of previous vascular organoid systems, enabling the generation of structurally and compositionally mature, adult-like arterial tissues.

Next, to evaluate whether the assembloid model recapitulates *in vivo* remodeling predicted by CausaLink, we compared ECM dynamics between MGL^high^ and MGL^low^ Mac assembloids and postnatal mouse arteries. In mouse arteries, ECM remodeling was characterized by an early peak in elastin deposition around P21, followed by a decrease accompanied by increased collagen and GAG content (**Fig. 1I**), reflecting a transition from elastin-driven elasticity toward collagen-mediated structural reinforcement. To experimentally validate these predictions, we assessed ECM remodeling dynamics in MGL^high^ and MGL^low^ Mac assembloids. In MGL^high^ Mac assembloids, elastin was enriched at the outer border on Day 3, prior to outer mesenchymal layer formation (**Fig. 5I, Extended Data Fig. 11**). During self-assembly, elastin expression increased within the outer layer by Day 5 but was reduced by Day 10. Conversely, elastin expression continued to increase through Day 10 in MGL^low^ Mac assembloids, reflecting dysregulated ECM dynamics. Similarly, collagen type IV expression gradually increased in the outer layer of MGL^low^ Mac assembloids, whereas it remained relatively low in MGL^high^ Mac assembloids through Day 10. Quantitative analyses confirmed these subset-specific remodeling patterns, revealing a marked reduction in elastin accompanied by an increase in collagen IV from Day 5 to Day 10 exclusively in MGL^high^ Mac assembloids, reflecting controlled ECM turnover associated with balanced arterial remodeling (**Fig. 5J**). Nevertheless, structural differences remain between assembloids and native arteries, including less organized ECM fiber alignment and density compared to the highly aligned collagen fiber alignment typically observed under physiological hemodynamic loading *in vivo*.

Collectively, these findings provide robust experimental support for our computational predictions, establishing *Mgl2* Macs as key causal regulators of balanced arterial growth and remodeling. These findings provide mechanistic insights into developmental vascular biology and highlights potential opportunities for tissue engineering and regenerative medicine through the integration of computational modeling with physiologically relevant human organoid systems.

## Discussion

Postnatal development of the proximal pulmonary artery begins with a rapid increase in blood flow but decrease in blood pressure that demands a unique evolving structure and function. Understanding this process requires integration of transcriptional, compositional, morphometric, and mechanical data to understand this complex biological process. Here, we introduce CausaLink, a causal inference framework designed to integrate multimodal datasets and uncover cross-scale causal relationships. CausaLink reconstructs directed gene–cell–trait causal hierarchies by learning a shared latent representation across independently acquired transcriptomic and biomechanical datasets, thereby overcoming the missing-data problem. Using this framework, we identified causal hierarchies linking *Mgl2^+^* Macs to balanced tissue-level remodeling in the developing pulmonary artery, providing mechanistic insight into postnatal vascular maturation.

CausaLink demonstrates strong potential for studying tissue-level remodeling processes involving interdependent traits—such as vessel diameter, wall thickness, and mechanical stiffness—that must evolve in a tightly coordinated manner. Disrupting this balance can lead to pathological outcomes including luminal encroachment, aneurysmal dilatation, or fibrosis. Traditional approaches including GWAS, TWAS, and eQTL analyses have linked genetic variants to individual traits^54–56^, but even their multivariate extensions remain largely static and associatve^14, 57^. They fail to capture causal hierarches or predict how genetic perturbations propagate through interrelated traits. CausaLink addresses this gap by coupling causal inference with dynamic, multi-trait simulation, thereby providing actionable multiscale insight into complex biological systems.

While this framework advances causal inference into the realm of predictive modeling, several areas offer opportunities for future development. First, although the framework integrates transcriptomic and biomechanical modalities, it does not yet leverage and integrate imaging data as direct inputs. Enabling end-to-end modeling with raw imaging data could expand CausaLink’s applicability to diverse biomedical datasets. Furthermore, while our current study focused on arterial tissue remodeling, future work should assess CausaLink’s generalizability to other remodeling contexts to fully evaluate its utility across biological systems.

The hiPSC-derived assembloid platform provided a crucial experimental system for validating CausaLink predictions. Incorporation of defined Mac subsets directly confirmed the causal role of MGL Macs in balanced arterial remodeling within a controlled, human-specific context. This humanized system thus enables precise mechanistic assessment of computational predictions. Although our findings identify *Mgl2*^+^ / MGL^high^ Macs as regulators of balanced arterial growth, further studies will be needed to define their developmental origin and distribution across vascular beds. Incorporating physiological hemodynamic stimuli—such as pulsatile blood flow or shear stress—could further improve the ability of the assembloid model to replicate native arterial dynamics and biomechanical signaling. However, the current assembloid lacks broader systemic physiological contexts, including endocrine signaling, neuronal innervation, and complex immune interactions inherent to living organisms, potentially affecting vascular remodeling outcomes. Validation in more physiologically integrated platforms, such as organ-on-chip platforms or more complex animal models including non-human primates, will therefore be critical to enhance the biological fidelity and generalizability. Future research should nevertheless include improving scalability, reproducibility, and standardization of assembloids to further advance the robust experimental platform, strengthening the connection between computational modeling and experimental validation from developmental biology and tissue remodeling toward effective clinical applications.

In conclusion, this work establishes a functional bridge between computational causal modeling and experimental validation at the tissue-systems level. We confirmed an MGL^high^ / *Mgl2*^+^ Mac subset as a critical regulator of balanced arterial growth and remodeling, uncovering a cell population and mechanism that had remained inaccessible to previous association-based analyses. These findings highlight the broader potential of causal modeling frameworks to not only map regulatory circuits but predict how genetic or cellular perturbations drive tissue-level outcomes —capabilities that traditional gene regulatory network models cannot achieve.

Within this framework, our study lays the foundation for a predictive approach to tissue engineering and regenerative medicine—one in which therapeutic outcomes across multiple traits are forecasted using mechanistically informed, multiscale causal maps, rather than through empirical trial-and-error.

## Acknowledgments

This work was funded by grants from Additional Ventures (AVCC to MED and JDH) and the American Heart Association Postdoctoral Fellowship (25POST1366397 to JH). This work was also supported by the Basic Science Research Program through the National Research Foundation of Korea (RS-2024-00411474 to HJP) and the Alchemist Project of the Korea Evaluation Institute of Industrial Technology (KEIT 20018560, NTIS 2410005252 to HJP) funded by the Ministry of Trade, Industry & Energy (MOTIE, Korea). This work was also supported by the VA VISN1 CDA (to EPM).

## Methods

### Animals

All animal experiments were approved by the Institutional Animal Care and Use Committee of Yale University. Healthy male C57BL/6J mice were studied at five ages: postnatal day 2 (P2, perinatal), P10 (neonatal), P21 (weaning/juvenile), P42 (sexual maturity), and P84 (adult). Following euthanasia, the right branch pulmonary artery (RPA) was isolated via a midline sternotomy and excised gently. Following ARRIVE Guidelines, excised vessels were randomly divided into three cohorts: those for biomechanical testing (n = 3-5) and histological analysis (n = 4-6) and those for those for RNA sequencing (n = 4-6 for bulk and n = 3 for single cell), each at P2, P10, P21, P42, P84, thus totaling 70+ mice for the study.

### Biomechanical phenotyping

We followed standard procedures for biomechanical phenotyping of murine vessels. Following euthanasia and excision of the RPA, the vessel was cleaned of excess perivascular tissue, cannulated on custom glass cannulae, and placed within a specimen bath containing a Hank’s buffered physiologic solution at room temperature to ensure a passive mechanical behavior. Following standard preconditioning, the vessels were subjected to a series of seven computer-controlled cyclic pressure-distension and axial force-extension protocols that generated pressure, diameter, axial force, and axial length data. The energetically favorable *in vivo* axial stretch was estimated as that value at which axial force changed little upon cyclic pressurization. Vessels were subjected to pressures up to which asymptotic behavior was observed in the pressure-diameter curves (20 mmHg at P2, 25 mmHg at P10, 30 mmHg at P21, and 40 mmHg at P42 and P84). The axial stretch was maintained constant at 95%, 100%, and 105% of the in vivo value. During the force-extension tests, vessels were subjected to axial forces up to the maximum value achieved during the 105% stretch pressure-diameter test while maintaining the luminal pressure fixed at four different values below the maximum testing pressure. Primary morphometric values were calculated at *in vivo* relevant loading conditions: luminal diameter from the measured outer diameter and wall thickness calculated by assuming transient incompressibility given measurements of unloaded cross-sections using a calibrated dissection microscope. Mural volume was estimated by assuming a cylindrical segment: the cross-sectional area multiplied by the vessel length (measured from the pulmonary valve to the first branch).

Mechanical quantities of interest such as circumferential and axial wall stretch, stress, and material stiffness as well as local distensibility were subsequently calculated from biaxial data and constitutive fits to the data at *in vivo* loading conditions. We used a 2-D formulation to quantify the mean mechanical behaviors of the vessels as good estimates of overall wall stress. Mean wall stresses in circumferential (θ) and axial (z) directions were determined experimentally as 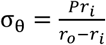 and 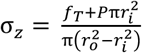, where *P* is the transmural pressure, *ri* internal radius, *ro* outer radius, and *fT* the transducer measured axial force. Under the assumption of incompressibility, inner radius 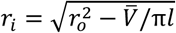, where *l* is the instantaneous length between the ligatures securing the vessel to the micropipettes and *V̅* is the volume of the wall in the unloaded state; *V̅* = π𝐿(𝑂𝐷^2^ − 𝐼𝐷^2^)/4 where *L* is the unloaded length, *OD* the unloaded outer diameter, and *ID =OD-2H,* the unloaded inner diameter. Mean biaxial wall stretches are λ_θ_ = (𝑟_𝑖_ + 𝑟_𝑜_)/(ρ_𝑖_ + ρ_𝑜_) and λ_𝑧_ = 𝑙/𝐿, where *ρi* and *ρo* are inner and outer radii in the unloaded configuration.

Following prior work, we further characterized the biaxial biomechanical data from the aforementioned seven cyclic testing protocols using a single nonlinear (pseudo)elastic stored energy function 𝑊 that has successfully described passive biaxial behaviors of proximal pulmonary arteries, namely 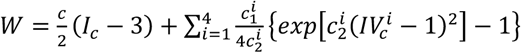, where 𝑐 (kPa), 𝑐_1_^𝑖^ (kPa) and 𝑐_1_^𝑖^ (-) are material parameters; 𝑖=1,2,3,4 represent four collagen-dominated families of fibers along axial, circumferential, and two symmetric diagonal directions, respectively. 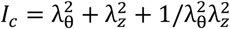 is the first invariant of the right Cauchy-Green tensor and 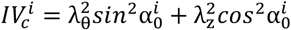 is the square of the stretch of the 𝑖*^th^* fiber family where α_0_^𝑖^ is the fiber angle relative to the axial direction in the reference configuration. Best-fit values of the material parameters and fiber angle were determined via nonlinear regression of biaxial data from all seven passive protocols (∼2800 data points per vessel). More details on parameter estimation can be found elsewhere, noting that the associated biaxial wall stress and material stiffness can be computed from first and second derivatives of 𝑊 with respect to an appropriate deformation metric.

### Histology

Following biomechanical testing, the samples were fixed in 10% neutral buffered formalin and stored in 70% ethanol at 4°C. Samples were embedded in paraffin, sectioned, and stained with Movat pentachrome (MOVAT) by a Yale histology core. Under bright-field illumination, MOVAT reveals elastin as black, collagen as grey-yellow, glycosaminoglycans as blue, cytoplasm as pink, and, if present, fibrin as dark red. Sections were imaged with an Olympus BX/51 microscope and an Olympus DP70 digital camera at 40X magnification. Complete cross-sections were obtained by stitching together sub-images with the Image Composite Editor software (Microsoft Research). The stitched images were subsequently analyzed using custom MATLAB scripts (https://github.com/yale-humphrey-lab/histology-analysis). that included background subtraction and pixel-based HSL-based thresholding. Three sections (technical replicates) were analyzed per vessel (five biological replicates), thus yielding 15 sections per age. Area fractions for elastin and cytoplasm were computed as the ratio of pixels corresponding to a stain divided by the total number of pixels in the image. The remaining pixels not categorized as elastin or cytoplasm in the histological sections are considered to be a combination of collagen and GAGs, with the area fraction of this collagen aggregate computed as 1 - area fraction of elastin plus cytoplasm^58^. Finally, additional unloaded morphometric measurements included luminal diameter (measured across the midpoint of the RPA cross-section) and wall thickness (measured as the radial distance between the endothelium and external adventitial boundary, averaged over four quadrants per section). Adventitial thickness was quantified specifically as the distance from the external elastic lamina to the external adventitial boundary.

### Bulk RNA Sequencing Analysis

Immediately upon euthanasia, RNA was isolated from the extralobar pulmonary arteries using an RNeasyMini Kit (Qiagen) according to the manufacturer’s specifications. Quality control measures ensured that samples were not degraded, with RNA integrity (RIN) > 7. RNA libraries were prepared with polyA selection. Whole-transcriptome sequencing was performed using a NovaSeq 6000 System (Illumina, Inc.) by The Yale Center for Genome Analysis. Sequenced reads were imported into CLC Genomics Workbench V23 (Qiagen) and, following quality control, reads were trimmed and aligned/mapped to a *Mus musculus* reference genome. Reads were then automatically processed using log counts per million (CPM) transformation with trimmed mean of M (TMM) adjustment.

### Single-Cell RNA Sequencing Analysis

#### Tissue dissociation

Following euthanasia, the excised extralobar pulmonary arteries were chopped mechanically and placed in 1.5 mg/ml collagenase (Gibco), 3 U/ml elastase (Worthington), 1 mg/ml DNASE I (Roche), 1.5 mg/ml dispase (Sigma-Aldrich), and 1 mg hyaluronidase (Sigma-Aldrich).

#### Library preparation and sequencing

Single-cell capture and library preparation were performed using the Chromium Single Cell 3′ v3.1 platform (10x Genomics) following the manufacturer’s protocol. Approximately 10,000 cells were loaded per sample to target ∼5,000 captured cells. After barcoding and cDNA amplification, sequencing libraries were constructed and sequenced on an Illumina NovaSeq 6000 system to a depth of ∼50,000 reads per cell. Cell Ranger software (10x Genomics, v6.1.1) was used to demultiplex raw reads, align to the mouse reference genome (mm10), and generate gene expression count matrices for each sample.

#### Pre-processing and cell clustering

Downstream analysis was carried out in R (v4.1) using the Seurat^59^ single-cell analysis or Loupe browser (v.7.0.1). Cells with fewer than 200 or greater than 20,000 detected UMIs or fewer than 300 detected features, or >20% mitochondrial gene content were excluded to remove low-quality captures. Datasets from all time points were normalized by SCTransform function and integrated using Seurat’s integration workflow to correct for batch effects across ages. Highly variable genes were identified and used for principal component analysis (PCA). The top 10 principal components were used to construct a shared nearest-neighbor graph, and unsupervised clustering was performed. A two-dimensional UMAP was computed for visualization of the cellular landscape across development (minimum distance: 0.5, number of neighbors: 500). Clustering analysis was performed with a resolution parameter of 0.4. Markers for each cluster were found using the FindAllMarkers function with a logFC threshold of 0.25 and a return threshold of 0.05. Clusters were identified by cross-referencing their highest expressed markers to the CellMarker (v 2.0)^60^, PanglaoDB (https://panglaodb.se/)^61^, and CZ CELLxGENE (https://cellxgene.cziscience.com/) databases. When necessary, we also searched markers through a manual literature search. For heatmaps, the integrated, scaled, and normalized dataset was down sampled to 100 cells and the Seurat package’s DoHeatmap function was used to plot the differentially expressed genes in each cluster.

#### Differentially expressed genes (DEGs)

To assess transcriptional changes during postnatal development of the pulmonary artery, we contrasted gene expression between annotated neonatal or adult SMC, FB, and Mac populations using Seurat’s FindMarkers function (Wilcoxon test, adjusted *P* < 0.05, log2FC > 0.25). Results were visualized using EnhancedVolcano package in R, highlighting significantly altered genes. For functional interpretation, top genes (ranked by log2FC) were analyzed via the Reactome Pathway Database, with enriched pathways (*FDR* < 0.05) used to identify lineage-specific programs involved in pulmonary artery maturation.

#### Cell-cell Interaction Inference

CellChat (v2.0)^31^ was used to infer intercellular communication networks across postnatal pulmonary artery development (P2, P10, P21, P42, P84). Seurat objects from each age were converted into CellChat objects, and the built-in CellChatDB ligand–receptor interaction database (∼2,000 pairs) was used for reference. Overexpressed ligands and receptors were identified per cluster, and pathway-level communication probabilities were computed using CellChat’s computeCommunProb and computeCommunProbPathway, with significance threshold set at *P* < 0.05. To visualize global signaling patterns, CellChat’s netVisual_circle and netVisual_heatmap were used. We quantified outgoing (sender) and incoming (receiver) signal strength for each cell type using CellChat’s netAnalysis_signalingRole, enabling identification of dominant signaling sources and targets at each developmental stage. This allowed us to track temporal shifts in cell–cell communication dynamics and nominate potential orchestration hubs during vascular maturation.

### CausaLink framework

#### Statistical modeling

Pairwise Pearson correlation coefficients were computed across 14 mechanical parameters to generate a correlation matrix, from which edges with absolute values ≥ 0.77 were retained to construct a mechanical property network. The graph was clustered using K-means (k = 2), and visualized via a spring layout in NetworkX, where node size reflected connectivity and color denoted cluster identity. Circumferential stiffness emerged as a central hub parameter based on clustering topology.

To identify genes associated with circumferential stiffness, a bulk RNA-seq dataset (n = 27) spanning the five developmental ages was integrated with group-level mean stiffness values using Partial Least Squares (PLS) regression. The model was implemented using the PLSRegression class from Scikit-learn (components = 10) and evaluated via fivefold repeated cross-validation (RepeatedKFold), with model fit assessed by R² and mean squared error metrics. Variable importance in projection (VIP) scores were computed using the decomposition of scores (t), weights (w), and loadings (q) as follows:

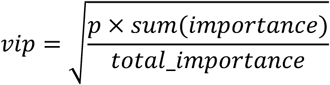

where importance is the diagonal of the product of t, q, and qᵀ, and *p* is the total number of predictors. Genes were ranked by VIP score, and the top 30 were selected for downstream analysis.

#### Cross-Modal Data Integration via Autoencoder

To integrate unpaired top15 screened genes and 4 key mechanical measurements across postnatal pulmonary artery development (P2, P10, P21, P42, P84), we implemented a custom cross-modal autoencoder framework in TensorFlow (v2.13). This model learns a shared latent representation between two modalities—gene expression (Modality A) and tissue mechanical data (Modality B)—without requiring one-to-one sample pairing.

Each modality was independently encoded into a shared latent space via dedicated encoders, followed by symmetric decoders for reconstruction. Cross-modal translation was implemented by decoding the latent representation of one modality into the feature space of the other. Training was conducted using a composite loss function comprising: (i) within-modality reconstruction loss (mean squared error), (ii) cycle-consistency loss to regularize cross-domain translation, and (iii) maximum mean discrepancy (MMD) loss to align latent distributions across domains. The model was trained for 1,000 epochs with a batch size of 16 and Adam optimizer (learning rate = 0.005), using randomly sampled mini-batches from each modality per step.

Post-training, we extracted latent representations from both domains and visualized them using UMAP (n_neighbors = 15, min_dist = 0.1). Feature importance was assessed by averaging the absolute weights of the first encoder layer and final decoder layer per modality. To evaluate biological plausibility, gene-to-mechanics translation was performed by passing gene-derived latent representations through the mechanical decoder. Predicted mechanical values were inverse-transformed and validated against empirical ranges. For each mechanical feature and age group, we quantified the proportion of translated values falling within the observed range and visualized the results with overlaid boxplots.

Latent embeddings from both modalities were co-embedded using UMAP, colored by age group and domain, and plotted to assess integration quality. Separate plots were generated before and after training to visualize improvements in modality alignment. Age-specific and modality-specific structure was preserved while achieving inter-domain mixing, demonstrating successful alignment of unpaired data across biological and physical feature spaces.

#### Causal gene-trait network discovery

To uncover directed causal relationships among prioritized genes and mechanical traits, a score-based causal discovery framework was applied to a matrix of 15 top-ranked genes (VIP-selected) across 44 integrated datasets. The DAS algorithm, implemented via the dodiscover package (v0.0), was used to infer a DAG. A contextual graph was first initialized using make_context(), and variable ordering was estimated using topological sorting informed by conditional independence scores. Resulting graphs—both the pruned causal DAG and the full order graph—were visualized using a customized graphviz-based pipeline.

#### In Silico Causal Effect Simulation

A probabilistic causal model was constructed using the ProbabilisticCausalModel class from the *DoWhy-GCM* library, based on a directed acyclic graph (DAG) inferred via the DAS algorithm. Causal mechanisms were defined using EmpiricalDistribution for root nodes and AdditiveNoiseModel for non-root nodes, with regression functions implemented via SklearnRegressionModel using either XGBRegressor (objective = “reg:squarederror”) or Lasso (alpha = 1.0). The model was trained on a gene expression matrix comprising 70 genes across 44 postnatal samples (P2–P84).

Gene perturbation simulations were performed by scaling the expression of *Fbln5* using fold-change multipliers calculated from a bulk RNA-seq dataset. For each multiplier, 1,000 interventional samples were generated using interventional_samples, and the mean values of four mechanical traits—diameter, wall thickness, volume, and circumferential stiffness—were computed. Confidence intervals for each estimate were obtained using the confidence_intervals function with 100 bootstrap resamples.

Multipliers and trait values were aligned to five postnatal timepoints (P2, P10, P21, P42, P84). Simulation results and corresponding empirical measurements were visualized using matplotlib.

### Experimental validation on Mac-arterial assembloids

#### Differentiation of THP-1-derived Macs

Human THP-1 monocytes were cultured in RPMI-1640 medium (Cytiva, Marlborough, MA, USA) supplemented with 10% fetal bovine serum (FBS; Gibco, Waltham, MA, USA). For differentiation into M0 Macs, cells were treated with 100 ng/mL phorbol 12-myristate 13- acetate (PMA; Sigma, St. Louis, MO, USA) for 24 hours. M0 Macs were further polarized into MGL^low^ Macs by treatment with 20 ng/mL interferon-γ (IFN-γ; Peptrotech, Cranbury, NJ, USA) and 1 mg/mL lipopolysaccharide (LPS; Sigma) for 3 days, or into MGL^high^ Macs by incubation with 20 ng/mL interleukin-10 (IL-10, Peprotech) and 20 ng/mL transforming growth factor-β1 (TGF-β1; Peprotech) for 3 days. All differentiation and polarization steps were conducted in RPMI-1640 medium supplemented with 10% FBS.

#### Flow cytometry analysis

THP-1 cells were seeded at a density of 3 × 10⁵ cells/well into 12-well plates and polarized as described above. Following polarization, cells were detached with 0.25% trypsin (Cytiva) and harvested. Cells were then fixed with 10% formalin for 10 min at room temperature, stained with APC-conjugated anti-CD301 (MGL) antibody (1:50; Miltenyi Biotec, Bergisch Gladbach, Germany) for 2 hours, and analyzed using a NovoCyte flow cytometer (Agilent, Santa Clara, CA, USA). Data analysis was performed by manual gating using NovoExpress software (version 1.6.2, Agilent).

#### Generation of Mac-arterial assembloids

BVOs were generated from hiPSCs (KSCBi005-A, National Stem Cell Bank of Korea) as previously described^62^. For BVO differentiation, hiPSC aggregates were formed by seeding single cells which were dissociated using 0.1% Accutase (Invitrogen, Waltham, MA, USA) into low-attachment plates in aggregation medium (KnockOut DMEM/F-12 medium (Gibco) supplemented with 20% KnockOut Serum Replacement (KSR, Gibco) and 1% Non-essential amino acids (NEAA, Gibco)) containing 50 mM Y-27632 (Tocris, Bristol, UK). Cell aggregates were induced into mesodermal differentiation by N2B27 medium (a 1:1 mixture of DMEM/F-12 and Neurobasal medium (Gibco) supplemented with B27 supplement (Gibco), and N2 supplement (Gibco)) containing 12 mM CHIR99021 (Tocris) and 30 ng/mL bone morphogenetic protein-4 (BMP-4; Peptrotech) for 3 days. For vascular specification of cell aggregates, medium was switched to containing 100 ng/mL vascular endothelial growth factor-165 (VEGF-165; Peprotech) and 2 mM forskolin (STEMCELL Technologies, Vancouver, Canada) and cultured for 2 days. On Day -6, cell aggregates were embedded in a collagen-Matrigel matrix composed of 3 mL of collagen type I solution (450 mL 0.1N NaOH, 187 mL 10X DMEM (Gibco), 37.8 mL HEPES (Gibco), 29.4 mL 7.5% sodium bicarbonate, 19.2 mL Glutamax (Gibco), 276 mL Ham’s F-12 (Gibco), 2 mL collagen type I (Corning, Corning, NY, USA)) and 1 mL growth factor reduced-Matrigel (Gibco). After the gelation of matrix containing differentiated cell aggregates, BVO culture medium (StemPro-34 SFM medium (Gibco) supplemented with 15% FBS, 100 ng/mL VEGF-165, and 100 ng/mL fibroblast growth factor-2 (FGF-2; R&D Systems, Minneapolis, MN, USA)) was added for vessel sprouting. On Day -1, embedded BVOs were manually harvested using 30- gauge needles and transferred into new low-attachment plates for overnight culture. The following day, individual BVOs were transferred into U-bottom low-attachment 96-well plates (one organoid/well) and co-cultured with THP-1-derived MGL^low^ or MGL^high^ Macs (1.5 × 10^4^ cells/well) in 100 mL of assembloid medium (a 1:1 mixture of Macs differentiation medium and BVO culture medium). Day 0 was defined as one day after the sprouting phase, at which point ECs expressing tip-cell markers emerged at the angiogenic front and mesodermal cells began sprouting from the aggregates. At this stage, vascular networks began to self-assemble into hBVOs within ultra-low attachment U-bottom wells, forming microvessel lumens lined by ECs surrounded by mural cells^63^. On day 5 of co-culture, an additional 100 mL of fresh assembloid medium was added to each well.

#### Cell proliferation assay of Macs in assembloids

THP-1-derived MGL^low^ or MGL^high^ Macs were labelled with carboxyfluorescein diacetate succinimidyl ester (CFDA-SE; Invitrogen) according to the manufacturer’s instructions prior to co-culture. Briefly, Macs were incubated with 5 mM CFDA-SE dye for 30 minutes at 37°C and subsequently washed twice with RPMI-1640 medium supplemented with 10% FBS. CFDA-SE-labelled Macs were then co-cultured with BVOs in 96-well plates to generate assembloids. At Day 5 or Day 10 of co-culture, assembloids were harvested, washed twice with PBS to remove non-migrated Macs, and dissociated into single cells by incubation with 0.1% Accutase for 30 minutes at 37°C. The dissociated cells were filtered through a 40-mM cell strainer, fixed in 10% formaldehyde and analyzed for CFDA-SE fluorescence intensity using NovoCyte flow cytometer. Flow cytometry data were analyzed using NovoExpress software.

#### Immunostaining

Assembloids were washed twice with PBS to remove non-migrated co-cultured Macs and then fixed in 10% formaldehyde overnight at 4°C. For immunohistochemistry staining of cryosection, fixed assembloids were embedded in 30% sucrose solution for 24 hours and transferred in gelatin solution within plastic cryomolds. Cryosections were blocked for 1 hour at room temperature with blocking buffer (PBS containing 3% FBS, 1% BSA (GenDepot, Katy, TX, USA), 0.5% Triton X-100 (Sigma), 0.5% Tween-20 (Amresco, Solon, OH, USA), and 0.01% (w/v) sodium deoxycholate), and then incubated with primary antibodies (**Extended Data Table 1**) diluted in blocking buffer overnight at 4°C. After washing with PBS-T (PBS containing 0.05% Tween-20) three times, cryosections were incubated with secondary antibodies diluted 1:1000 in blocking buffer for an hour at room temperature. Nuclei were counterstained with DAPI (Invitrogen, 1:1000 dilution in PBS) for 10 minutes, followed by two washes with PBS-T. Samples were mounted using Fluoroshield (Sigma). For whole-mount immunostaining, fixed assembloids were blocked with blocking buffer for 2 hours at room temperature and incubated with primary antibodies (**Extended Data Table 1**) overnight at 4°C on a rocking shaker. After washing with PBS-T on an orbital shaker, samples were incubated with secondary antibodies diluted 1:250 in blocking buffer for 2 hours at room temperature. Whole assembloids were counterstained with DAPI and mounted onto 22 ⨉60 mm^2^ coverslips using Fluoroshield.

#### Immunofluorescence confocal imaging

Immunostained assembloid sections or whole-mounted samples were imaged using confocal microscopes (Eclipse TE200, Nikon, Tokyo, Japan; Stellaris 5, Leica Microsystems, Wetzlar, Germany) equipped with 10’ and 20’ objectives. Z-stack images were acquired to reconstruct the three-dimensional structures of assembloids and processed using LAS X software (Leica Microsystems).

#### Image and statistical analysis

Confocal images were processed and analyzed using ImageJ software (NIH, Bethesda, MD, USA). Vessel diameter (CD31^+^ structures) and thickness of outer layer were manually measured from orthogonal views of Z-stack images of assembloids. Statistical analyses were performed using GraphPad Prism software (GraphPad Software, Boston, MA, USA). Data are presented as mean ± SEM unless otherwise stated. The specific statistical tests used are described in the corresponding figure legends.

## Data and Code Availability

All data supporting the findings of this study are available within the paper and its Supplementary Information. Custom code for the cross-modal integration and causal discovery is available from the corresponding author upon reasonable request.

