## Supplemental data for "A Cross-scale Causal Mapping Framework Pinpoints Macrophage Orchestrators of Balanced Arterial Development"

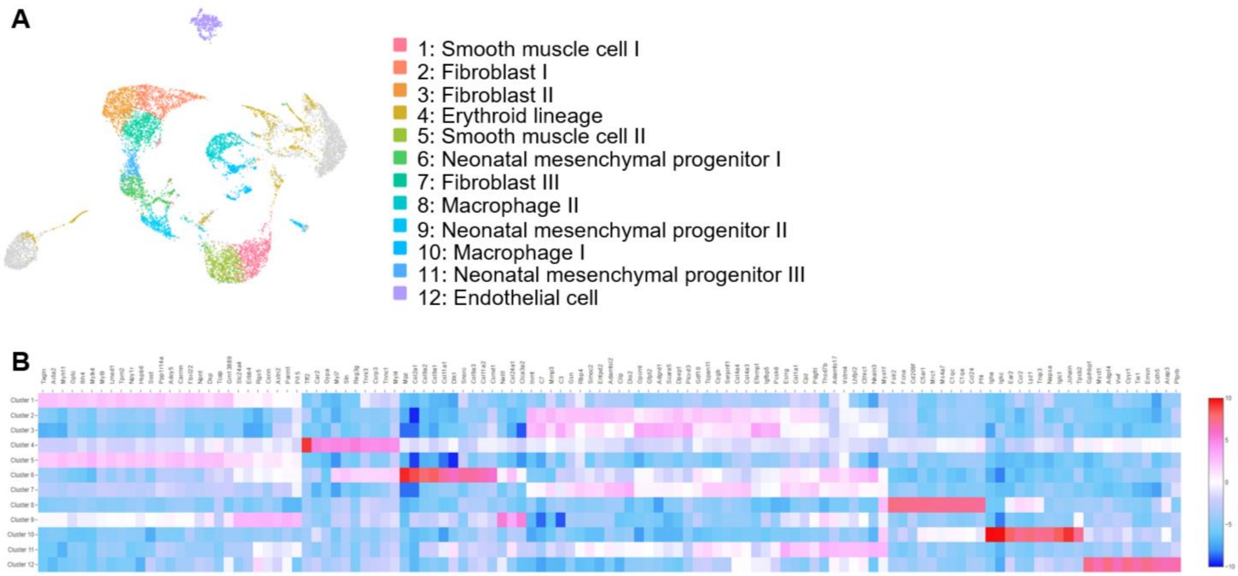


**Extended Data Fig. 1 Single-cell transcriptomic landscape of proximal pulmonary artery.** (**A**) UMAP projection of single-cell transcriptomes from postnatal pulmonary artery tissue showing 12 transcriptionally distinct cell clusters. Colors indicate unsupervised clusters, including FBs (clusters 2, 3, 6), SMCs(clusters 1, 5), neonatal mesenchymal progenitors (clusters 7, 9, 11), macrophages (clusters 8, 10), endothelial cells (cluster 12), and erythroid lineage cells (cluster 4). (**B**) Heatmap of top marker genes for each cluster identified by differential expression analysis. For each cluster, the most distinctive genes are shown, highlighting transcriptional features that define lineage identity and state.

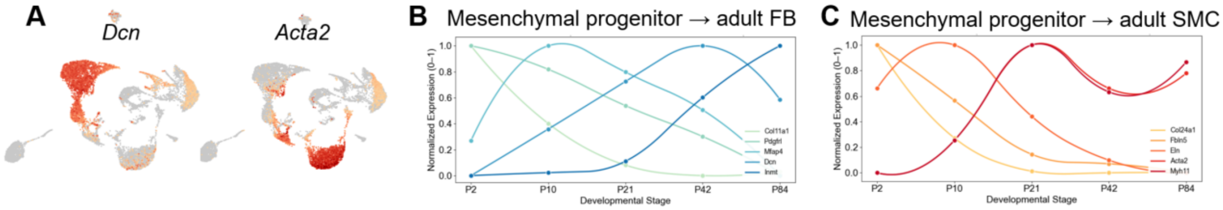


**Extended Data Fig 2. Time-course transcriptomic profiling of neonatal mesenchymal progenitors, FBs, and SMCs in developing mice pulmonary artery** (**A**) UMAP plots showing spatial projection of gene expression for selected FB (*Dcn*, left) and SMC–related (*Acta2*, right) markers across cells from postnatal pulmonary artery tissues. Warmer colors indicate higher gene expression. Temporal expression trends of (**B**) FB-associated genes (*Col1a1*, *Pdgfrl*, *Mfap4*, *Dcn*, *Inmt*) and (**C**) SMC–associated genes (*Col24a1*, *Eln*, *Fbln5*, *Acta2*, *Myh11*) normalized across five developmental stages (P2 to P84). Lines represent cubic spline interpolation of scaled average expression per stage. All expression values are min–max scaled (0–1) per gene.

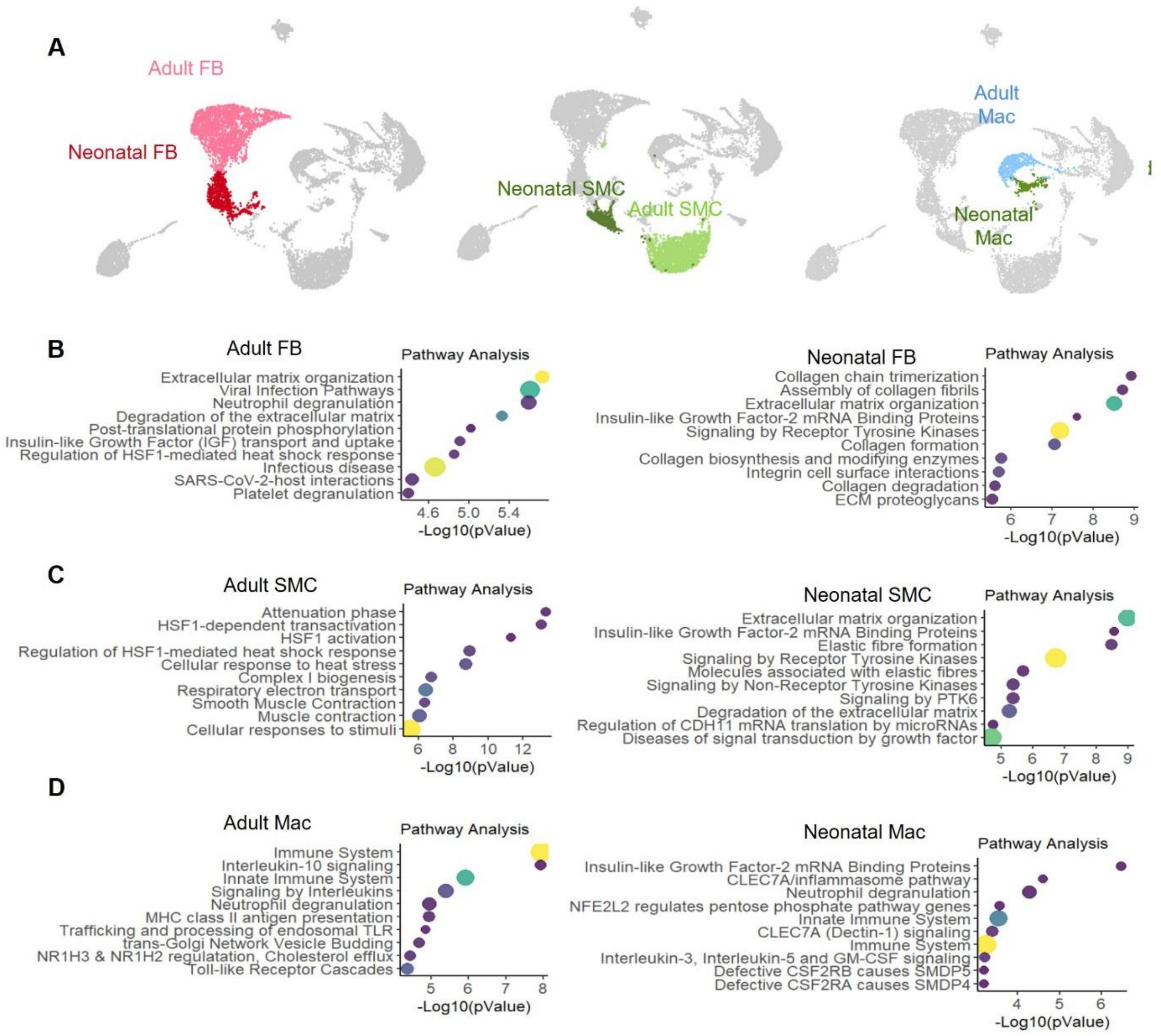


**Extended Data Fig 3. Neonatal (before P21) and adult (after P21) fibroblast, smooth muscle, and macrophage subsets exhibit distinct pathway enrichment signatures during pulmonary artery development.** (**A**) UMAP embedding of postnatal pulmonary artery single-cell transcriptomes, highlighting FB, SMC, and Mac stratified by developmental stage (neonatal vs. adult). Distinct spatial segregation is observed between neonatal and adult FB (left, red/pink), SMC (middle, green shades), and Mac (right, blue/cyan). (**B–D**), Pathway enrichment analysis comparing neonatal and adult subpopulations for each lineage: (**B**) FB, (**C**) SMC, and (**D**) Mac. Enrichment was performed using Reactome databases on differentially expressed genes. Dot size indicates gene ratio; color corresponds to significance (–$\log_{10}$(p-value)). Neonatal FB and SMC are enriched for matrix synthesis, collagen organization, and IGF-related pathways, while adult FB and SMC show enrichment in heat shock response and stress-related signaling. Adult Macs display immune-regulatory and antigen presentation pathways, contrasting with innate immune activation in adult Macs.

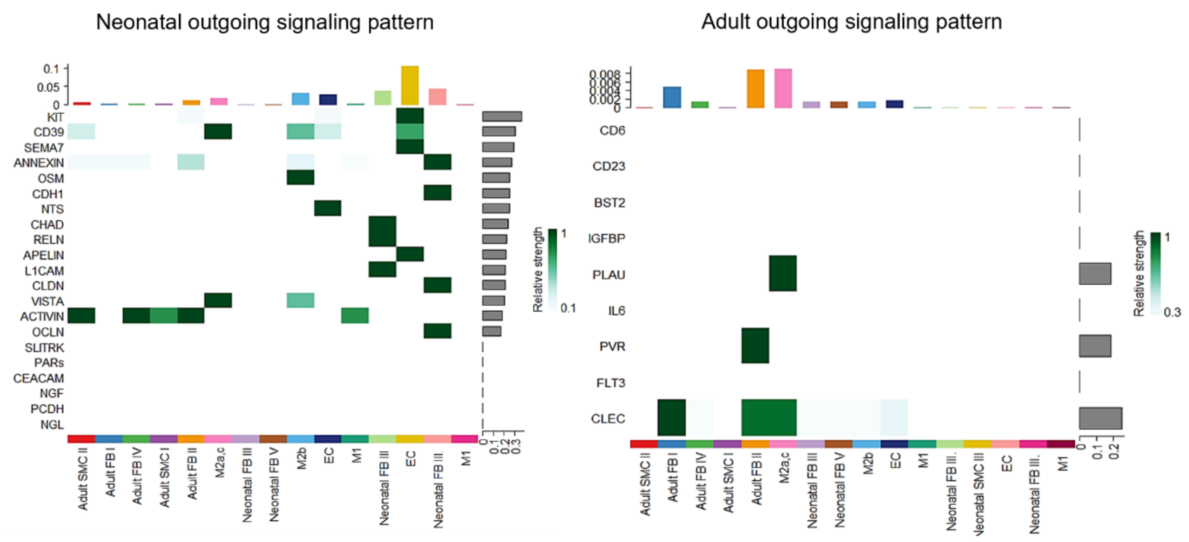


**Extended Data Fig 4. Stage-specific cell-cell interaction between neonatal and adult stages, visualizing changes in ligand-receptor interactions.**

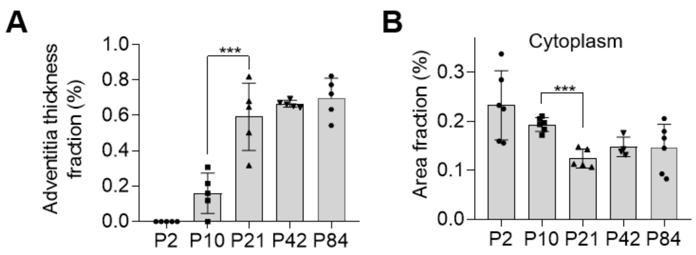


**Extended Data Fig 5. Tissue architecture changes during postnatal development.** (**A**) Quantification of adventitia thickness fraction, showing progressive remodeling (n = 5). (**B**) Quantification of cytoplasm area fractions across developmental stages. Data are presented as mean ± SEM. A one-way analysis of variance and Tukey's test were performed for statistical analysis (*P < 0.05, ***P < 0.001).

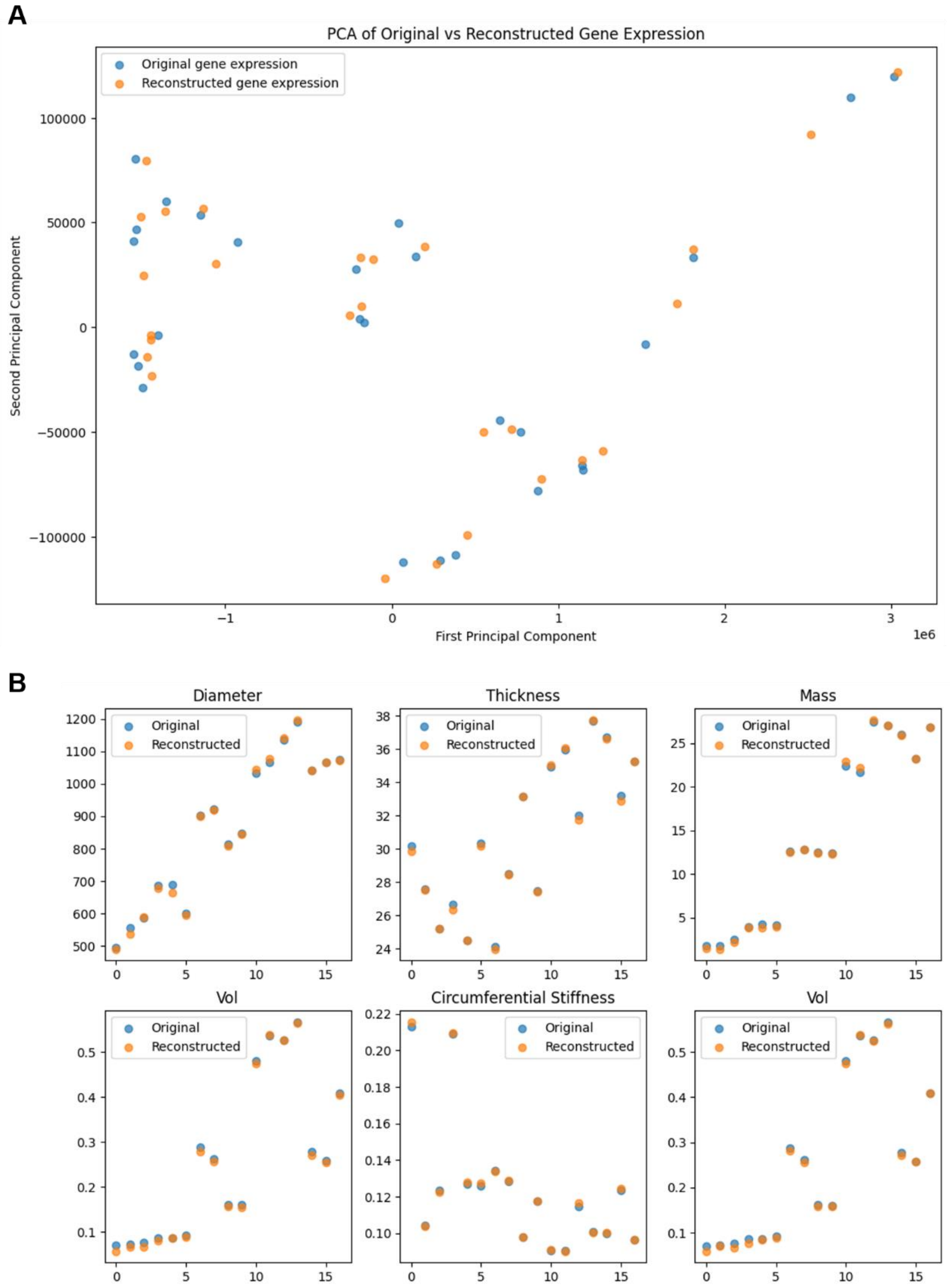


**Extended Data Fig 6. Validation of cross-modal autoencoder performance.** (**A**) PCA of original and reconstructed gene expression demonstrates close overlap, indicating preservation of variance structure. (**B**) Reconstructed biomechanical traits (diameter, thickness, mass, volume, circumferential stiffness) closely match original measurements across samples, confirming accurate cross-modal reconstruction.

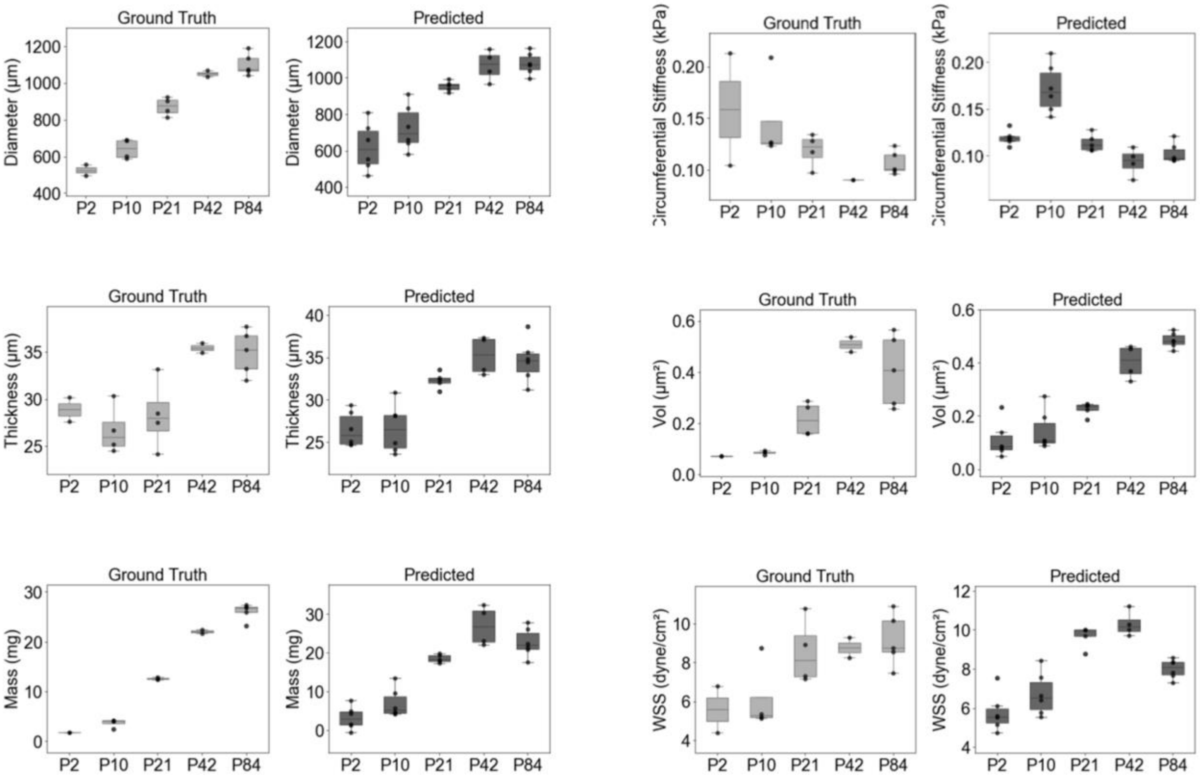


**Extended Data Fig 7. Cross-modal translation of gene expression data predicts tissue-scale structural and mechanical features of the postnatal pulmonary artery.** Box plots comparing measured (Ground Truth) and model-predicted (Predicted) values of six physiological properties across five postnatal developmental stages (P2, P10, P21, P42, P84): Diameter, Wall Thickness, Circumferential Stiffness, Wall Shear Stress (WSS), Mass, and Volume. The model accurately recapitulates the developmental trajectories of both structural (e.g., diameter, mass) and functional (e.g., stiffness, WSS) traits, capturing stage-specific growth and remodeling dynamics. Each box represents the interquartile range (IQR), with median lines and whiskers indicating 1.5× IQR; dots represent individual biological replicates. n =3-5 mice per stage.

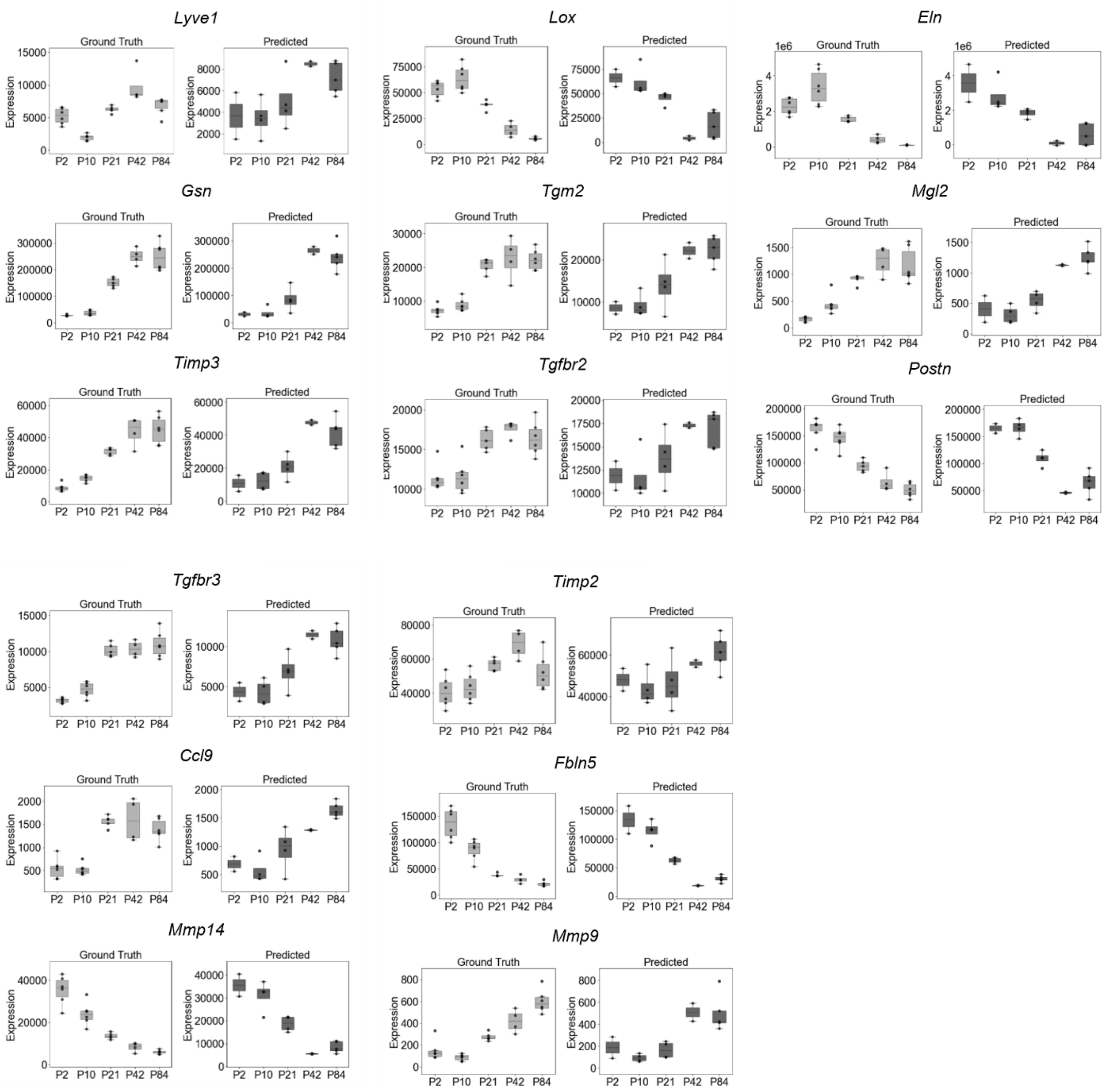


**Extended Data Fig 8. Cross-modal translation of biomechanical data predicts tissue-scale gene expression of the postnatal pulmonary artery.** Box plots comparing measured (Ground Truth) and model-predicted (Predicted) values of 15 gene expression across five postnatal developmental stages (P2, P10, P21, P42, P84). The model accurately recapitulates the developmental trajectories of both structural (e.g., diameter, mass) and functional (e.g., stiffness, WSS) traits, capturing stage-specific growth and remodeling dynamics. Each box represents the interquartile range (IQR), with median lines and whiskers indicating 1.5× IQR; dots represent individual biological replicates. n =3-5 mice per stage.

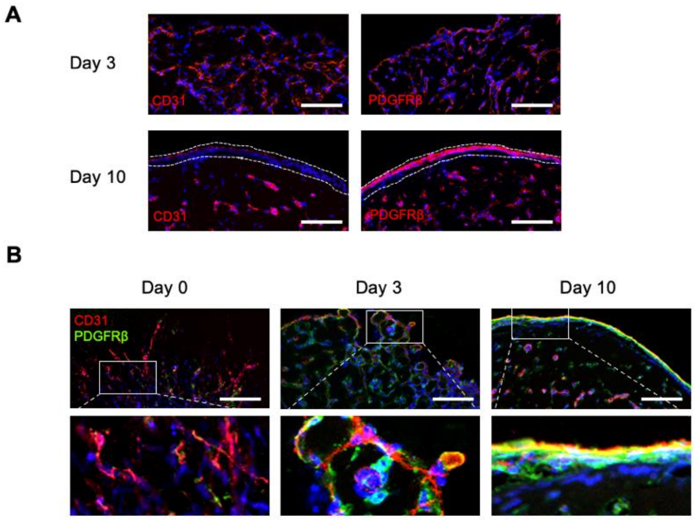


**Extended Data Fig 9. Structural organization of the outer region in hBVOs.** (A) Representative images showing the spatial arrangement of CD31$^{+}$ endothelial cells (ECs) and PDGFRβ$^{+}$ mesenchymal cells in the outer region of blood vessel organoids at Day 3 and Day 10. By Day 10, the CD31$^{+}$ EC layer localizes to the outermost boundary of outer layer (dotted line), with the PDGFRβ+ mesenchymal layer positioned directly beneath. Scale bars = 100 μm. (B) Temporal changes in the localization of CD31 and PDGFRβ expression in the outer region from Day 0 to Day 10, illustrating dynamic reorganization during self-assembly. Scale bars = 100 μm.

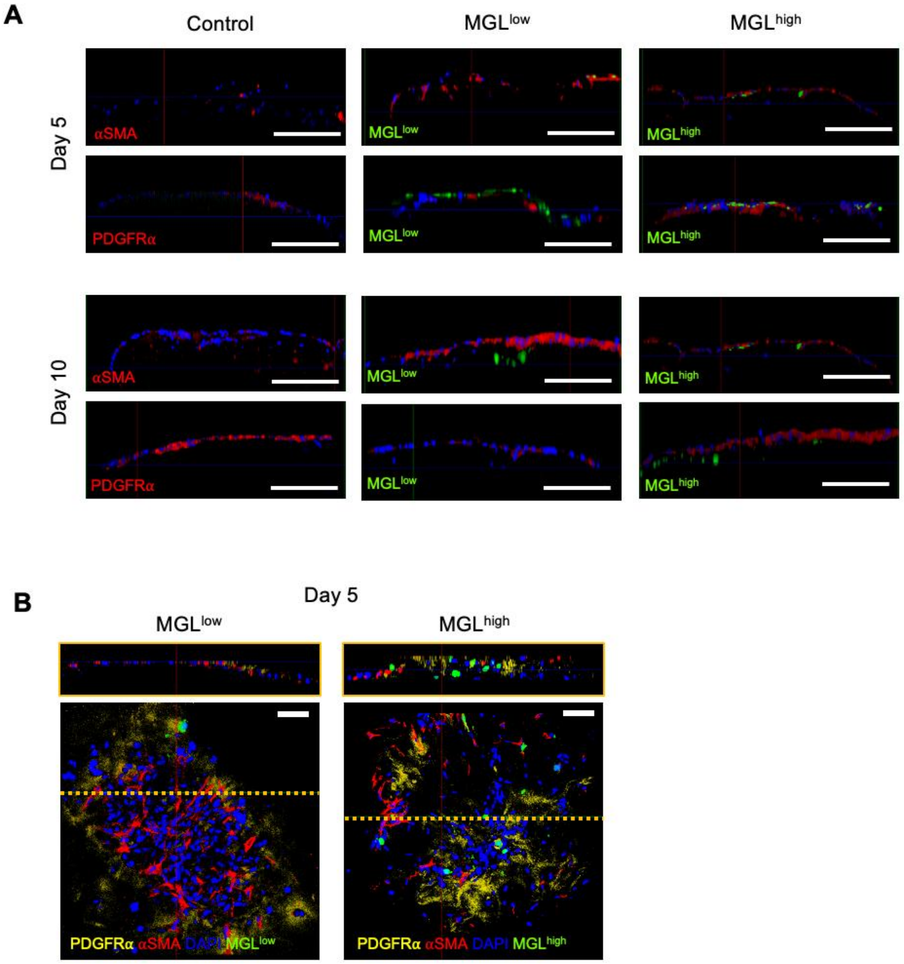


**Extended Data Fig 10. Macs-driven reorganization of FBs and SMCs in the mesenchymal layer of assembloids.** (A) Representative images showing dynamic changes in the outer mesenchymal layer of assembloids co-cultured with control, $MGL^{low}$, $MGL^{high}$ macrophages (Macs). $\alpha SMA^+$ smooth muscle cells (SMCs) and $PDGFR\alpha^+$ fibroblasts (FBs) form distinct but spatially reorganized layers depending on Mac subtype. (B) Spatial distribution of cell types in Day 5 assembloids co-cultured with $MGL^{low}$ or $MGL^{high}$ Macs. The frequency of αSMA and PDGFRα co-expressing cells is higher in the $MGL^{high}$ condition, with Macs frequently localized near these FB-SMC progenitor populations. Scale bars = 200 μm.

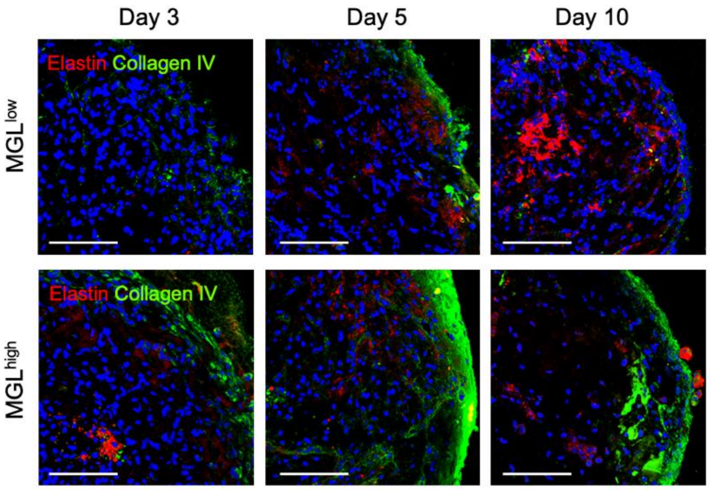


**Extended Data Fig 11. Mac-driven changes in ECM composition within arterial assembloids.** Elastin expression exhibited dynamic changes across both the inner and outer regions, showing early enrichment followed by a gradual decrease in $MGL^{high}$ assembloids. Collagen type IV expression was primarily localized to the outer region, peaking at Day 5. Scale bars = 50 μm.

| Name | Company | Catalog | Host animal | Dilution |
|---|---|---|---|---|
| APC-conjugated anti-CLEC10A | Miltenyi Biotec | Cat# 130-132-465 | Mouse | 1:500 |
| Anti-CD31 | Abcam (Cambridge, UK) | Cat# ab28364 | Rabbit | 1:1000 |
| Anti-αSMA | Abcam | Cat# ab7817 | Mouse | 1:1000 |
| Anti-PDGFRα | Santa Cruz Biotechnology (Dallas, TX, USA) | Cat# sc-21789 | Mouse | 1:500 |
| Anti-PDGFRα | Abcam | Cat# ab203491 | Rabbit | 1:500 |
| Anti-Collagen type IV | Abcam | Cat# ab6586 | Rabbit | 1:500 |
| Anti-PDGFRβ | R&D systems | Cat# MAB1263 | Mouse | 1:1000 |
| Anti-Elastin | Invitrogen | Cat#MA1-27129 | Mouse | 1:500 |

**Extended Data Table 1**. **Antibody usage**
